# tascCODA: Bayesian tree-aggregated analysis of compositional amplicon and single-cell data

**DOI:** 10.1101/2021.09.06.459120

**Authors:** Johannes Ostner, Salomé Carcy, Christian L. Müller

## Abstract

Accurate generative statistical modeling of count data is of critical relevance for the analysis of biological datasets from high-throughput sequencing technologies. Important instances include the modeling of microbiome compositions from amplicon sequencing surveys and the analysis of cell type compositions derived from single-cell RNA sequencing. Microbial and cell type abundance data share remarkably similar statistical features, including their inherent compositionality and a natural hierarchical ordering of the individual components from taxonomic or cell lineage tree information, respectively. To this end, we introduce a Bayesian model for **t**ree-aggregated **a**mplicon and **s**ingle-**c**ell **co**mpositional **d**ata **a**nalysis (tascCODA) that seamlessly integrates hierarchical information and experimental covariate data into the generative modeling of compositional count data. By combining latent parameters based on the tree structure with spike-and-slab Lasso penalization, tascCODA can determine covariate effects across different levels of the population hierarchy in a data-driven parsimonious way. In the context of differential abundance testing, we validate tascCODA’s excellent performance on a comprehensive set of synthetic benchmark scenarios. Our analyses on human single-cell RNA-seq data from ulcerative colitis patients and amplicon data from patients with irritable bowel syndrome, respectively, identified aggregated cell type and taxon compositional changes that were more predictive and parsimonious than those proposed by other schemes. We posit that tascCODA^1^ constitutes a valuable addition to the growing statistical toolbox for generative modeling and analysis of compositional changes in microbial or cell population data.

## 1 INTRODUCTION

Next-generation sequencing (NGS) technologies have fundamentally transformed our ability to quantitatively measure the molecular make-up of single cells (Shalek et al., 2013), tissues (Regev et al., 2017; Karlsson et al., 2021), organs (He et al., 2020), as well as microbiome compositions in and on the human body (Human Microbiome Project Consortium, 2012). Single-cell RNA sequencing (scRNA-seq) (Tang et al., 2009; Shalek et al., 2013; Macosko et al., 2015) has become the key technology for recording the transcriptional profiles of individual cells across different tissue types (Regev et al., 2017) and developmental stages (Griffiths et al., 2018), and for determining cell type states and overall cell type compositions (Trapnell, 2015). Cell type compositions provide informative and interpretable representations of the noisy high-dimensional scRNA-seq data and are typically derived from clustering characteristic gene expression patterns in each cell (Duo` et al., 2018; Traag et al., 2019), followed by analysis of the expression levels of marker genes (Luecken and Theis, 2019). As a by-product, these workflows also yield a hierarchical grouping of the cell types, either derived from the clustering procedure or determined by known cell lineage hierarchies. Determining changes in cell type populations across conditions can give valuable insight into the effects of drug treatment (Tsoucas et al., 2019) and disease status (Smillie et al., 2019), among others.

Complementary to scRNA-seq data collection, amplicon or marker-gene sequencing techniques provide abundance information of microbes across human body sites (Human Microbiome Project Consortium, 2012; Lloyd-Price et al., 2017; McDonald et al., 2018). Current estimates suggest that the human microbiome, i.e., the collection of microbes in and on the human body, outnumber an individual’s somatic and germ cells by a factor of 1.3-10 (Turnbaugh et al., 2007; Sender et al., 2016). Starting from the raw read counts, amplicon data are typically summarized in count abundance tables of operational taxonomic units (OTUs) at a fixed sequence similarity level or, alternatively, of denoised amplicon sequence variants (ASVs). The marker genes also allow taxonomic classification and phylogenetic tree estimation, thus inducing a hierarchical grouping of the taxa. To reduce the dimensionality of the data set and guard against noisy and low count measurements, the taxonomic grouping information is often used to aggregate the data at a fixed taxonomic rank, e.g., the genus or family rank. Shifts in the population structure of taxa have been implicated in the host’s health and have been associated with various diseases and symptoms, including immune-mediated diseases (Round and Palm, 2018), Crohn’s disease (Gevers et al., 2014), and Irritable Bowel Syndrome (IBS) (Ford et al., 2017).

In the present work, we exploit the remarkable similarities between scRNA-seq-derived cell type data and amplicon-based microbial count data and propose a statistical generative model that is applicable to both data modalities: the Bayesian model for **t**ree-aggregated **a**mplicon and **s**ingle-**c**ell **CO**mpositional **D**ata **A**nalysis, in short, tascCODA. Our model assumes that count data are available in the form of a *n × p*-dimensional count matrix *Y* containing the counts of *p* different cell types or microbial taxa in *n* samples, a covariate matrix *n × d*-dimensional *X* carrying metadata or covariate information for each sample, and a tree structure with *p* leaves that imposes a hierarchical order on the count data *Y*. Since both amplicon and scRNA-seq technologies are limited in the amount of material that can be processed in one sample, the total number of counts in rows of Y do not reflect total abundance measurements of the features but rather relate to the efficiency of the sequencing experiment itself (Gloor et al., 2017). This implies that the counts only carry relative abundance information, making them essentially compositional data (Aitchison, 1982).

tascCODA is a fully Bayesian model for tree-aggregated modeling of count data and is a natural extension of the scCODA model, recently introduced for compositional scRNA-seq data analysis (Büttner et al., 2020). At its core, tascCODA models the count data *Y* via a Dirichlet Multinomial distribution and associates count data and covariate information via a log-link function. To encourage sparsity in the underlying associations between the covariates and the hierarchically grouped features, tascCODA exploits recent ideas from tree-guided regularization and the spike-and-slab LASSO (Ročková and George (2018)). This allows tascCODA to perform tree-guided sparse regression on compositional responses with any type or number of covariates. In particular, in the presence of a single binary covariate, e.g., a condition indicator, tascCODA allows to perform Bayesian differential abundance testing. More generally, however, tascCODA enables to determine how host phenotype, such as disease status, host covariates such as age, gender, or an individual’s demographics, or environmental factors jointly influence the compositional counts. Finally, incorporating tree information into the inference allows tascCODA to not only identify associations between individual features, but also entire groups of features that form a subset of the tree.

tascCODA complements several recent statistical approaches, in particular, from the field of microbiome data analysis, some of which also use the concept of tree-guided models. Chen and Li (2013) were among the first to use the sparse Dirichlet-Multinomial model to connect compositional count data with covariate information in a penalized maximum-likelihood setting. Wadsworth et al. (2017) were the first to use a similar model in a Bayesian setting. Both adaANCOM (Zhou et al. (2021a)) and the Logstic-tree normal model (Wang et al. (2021)) use the Dirichlet-tree (multinomial) model (Wang and Zhao (2017)) to determine differential abundance of microbial taxa via a product of Dirichlet distributions at each split. These methods restrict themselves, however, to fully binary trees. One the other hand, the trac method (Bien et al., 2021)) uses tree-guided regularization (Yan and Bien, 2021)) in a maximum-likelihood-type framework to predict continuous outcomes from compositional microbiome data.

In its present form, the Bayesian model behind tascCODA is ideally suited for data sets of moderate dimensionality, typically *p* < 100, yet can handle extremely small sample sizes *n*. Since amplicon datasets are usually high-dimensional in the number of taxa and exhibit high overdispersion and excess number of zeros, we focus on the analysis of genus-level microbiome data. In the context of cell type compositional data, on the other hand, often only very few replicate samples are available (Büttner et al., 2020). Here, tascCODA can leverage well-calibrated prior information to operate in low-sample regimes where frequentist methods likely fail.

The remainder of the paper is structured as follows. In the next section, we introduce the tascCODA model and describe the computational implementation. In Section 3, we describe and discuss synthetic data benchmarks and provide two real-world applications, on human single-cell RNA-seq data from ulcerative colitis patients and amplicon data from patients with irritable bowel syndrome. Finally, we summarize the key points in Section 4 and present considerations about future extensions of the method. A flexible and user-friendly implementation of tascCODA is available in the Python package *tascCODA*^2^. All results in this paper are fully reproducible and available on Zenodo^3^.

## 2 MATERIALS AND METHODS

### 2.1 Model description

We start with formally describing the problem at hand. Let *Y* ∈ ℝ^*n*×*p*^ be a count matrix describing *n* samples from *p* features (e.g., cell types, microbial taxa, etc.), and *X* ∈ ℝ^*n*×*d*^ be a matrix that contains the values of *d* covariates of interest for each sample. Due to the technical limitations of the sampling procedure, the sum of counts in each sample, 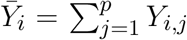 must be seen as a scaling factor, making the data compositional (Gloor et al. (2017)). Additionally, *Y* is hierarchically ordered by a multifurcating tree 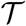 with *p* leaves and *t* internal nodes. Let *v* = *p* + *t* denote the total number of nodes in 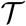. 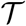 can be represented via a binary ancestor matrix *A* ∈ {0, 1}^*p*×*v*^:

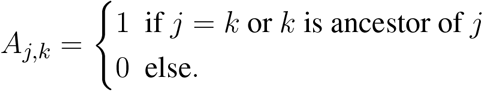

Our goal is to determine whether the abundance of single features (leaves of 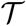) or entire subtrees are associated with the covariates in *X*. Hereby, a credibly changing subtree implies that the features contained in it are affected by the condition in the same manner (Figure 1A).

**Figure 1.**
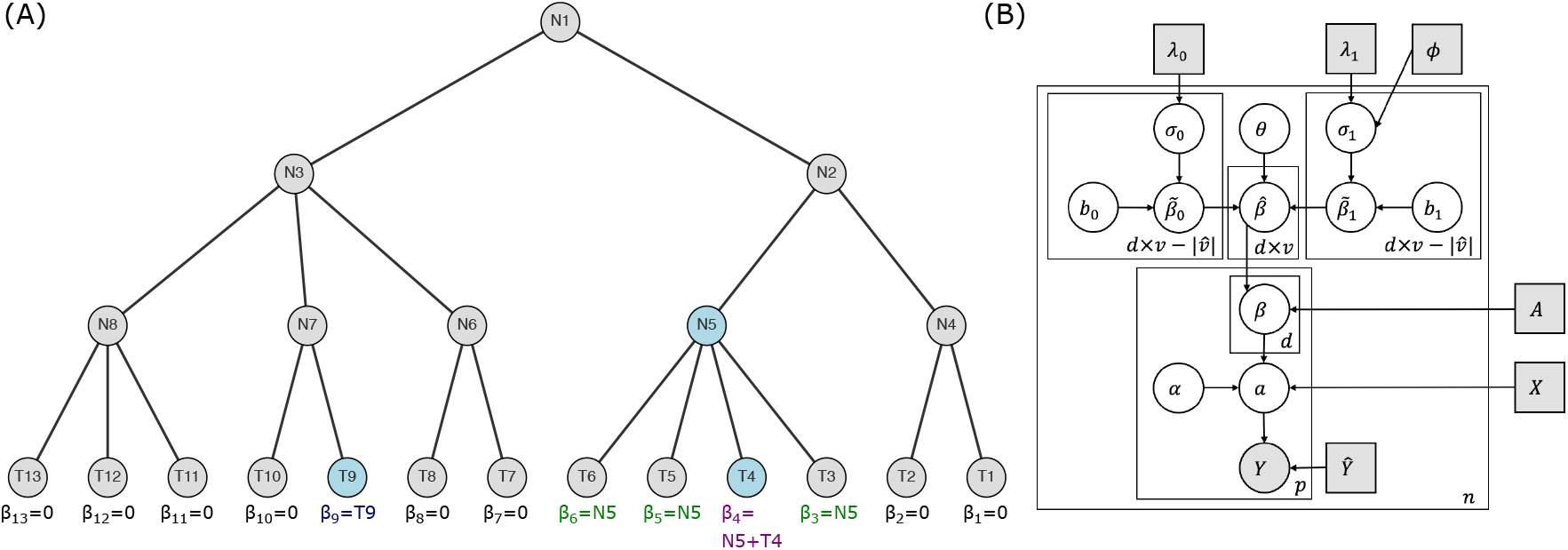
Intuition behind tascCODA, **(A)** a multifurcating tree structure 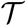 with internal nodes N1, … N8, and tips T1 … T13. If the blue nodes N5, T4, and T9 are assigned nonzero effects by tascCODA, the aggregated effects on the node level are displayed as *β*_1_ … *β*_13_ at the bottom. **(B)** Plate representation of the tascCODA model. Grey squares indicate fixed parameters and input variables that are either part of or directly calculated from the data. The grey circle represents the output count matrix, white circles show latent variables.

#### 2.1.1 Core model with tree aggregation

tascCODA posits a Dirichlet-Multinomial model for *Y_i_,*· for each sample *i* ∈ 1 … *n*, thus accounting for the compositional nature of the count data. The covariates are associated with the features through a log-linear relationship. We put uninformative Normal priors on the base composition *α*, which describes the data in the case *X_i_,*· = 0:

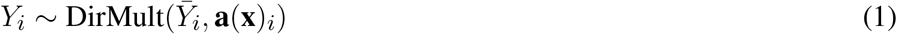

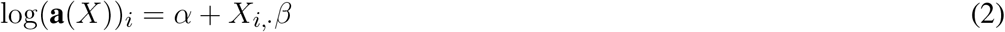

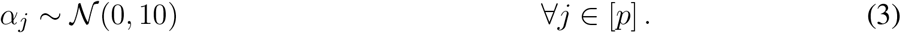

The total count 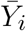 is directly inferred from the data for each sample. The effect of the *l*-th covariate on the *j*-th feature is therefore given by *β_l,j_*.

We now use a variant of the tree-based penalty formulation of Yan and Bien (2021) to model common effects at each internal node of 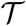 in addition to the effects on the leaves. We define a node effect matrix 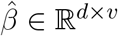 and calculate effects on the tips of the tree by multiplying with the ancestor matrix:

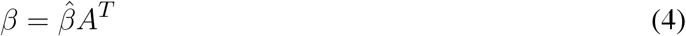

Thus e effect of covariate *l* on feature *k* is the sum over the effects of *l* on all ancestors of *k*, 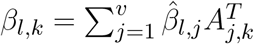. Figure 1A illustrates this tree-based aggregation process.

#### 2.1.2 Spike-and-slab lasso prior

To ease model interpretability, many statistical models provide a mechanism for sparsifying model parameters. In high-dimensional linear regression, this can be achieved via the lasso (Tibshirani, 1996), which adds an 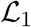-penalty on the regression coefficients. In Bayesian modeling, spike-and-slab priors are a popular choice to perform automatic model selection. Recently, (Ročková and George, 2018) developed a connection between the two approaches in the form of the spike-and-slab lasso prior, which provides a Bayesian equivalent to penalized likelihood estimation. Here, the effect of interest is described as a mixture of two double-exponential priors with different rates *λ*_0_, *λ*_1_ and a mixture coefficient *θ*:

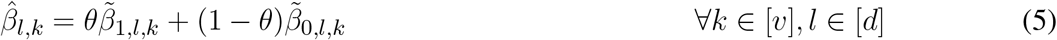

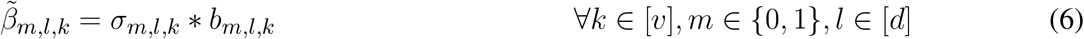

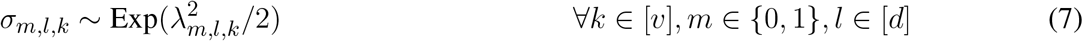

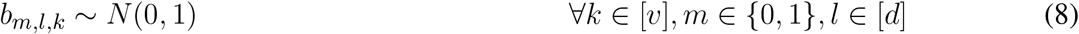

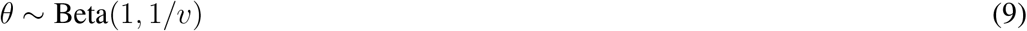

This prior can be reformulated as a likelihood penalty function that finds a balance between weak and strong penalization by *λ*_1_ and *λ*_0_, respectively (See Supplementary material section 1.2). As recommended by Ročková and George (2018), we use the non-separable version of the spike-and-slab lasso prior, which provides self-adaptivity of the sparsity level and an automatic control for multiplicity via a Beta prior on *θ* (Bai et al. (2020a); Scott and Berger (2010)). We further set *λ*_0*,l,k*_ = 50 ∀*k* to achieve a strong penalization in the “spike” part of the prior, leaving *λ*_1*,l,k*_ as our only parameter that controls the total amount of penalty applied at larger effect values.

#### 2.1.3 Node-adaptive penalization

We use a variant of the strategy proposed by Bien et al. (2021) to make the strength of the regularization penalty dependent on the corresponding node’s position in the tree. We introduce the following sigmoidal scaling:

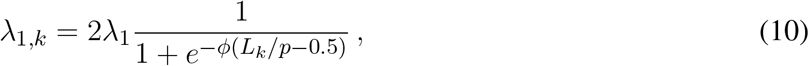

where *λ*_1_ = 5 is the default value for the penalty strength, *L_k_* is the number of leaves that are contained in the subtree of node *k*, and *φ* acts as a scaling factor based on the tree structure. If *φ* = 0, the default in tascCODA, all nodes are penalized equally with *λ*_1_, while for *φ* < 0, effects on nodes with larger subtrees, located closer to the root of the tree, are penalized less and are therefore more likely to be included in the model. If *φ* > 0, a solution that comprises more diverse effects on leaf nodes will be preferred. Thus, the parameter *φ* provides a way to trade off model accuracy with the level of aggregation. We discuss the behavior of the spike-and-slab LASSO penalty and the choice of *λ*_0,1_ in more detail in the Supplementary material.

#### 2.1.4 Reference feature

Since the data at hand is compositional, model uniqueness and interpretability are only guaranteed with respect to a reference. Popular choices include picking one of the *p* features or the (geometric) mean over multiple or all groups (Fernandes et al., 2014). Following the scCODA model, we pick a single reference feature prior to analysis (Büttner et al., 2020). Technically, this is achieved by choosing one feature 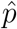 that is set to be unchanged by all covariates. Let 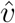 be the set of ancestors of 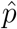. By forcing 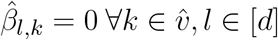, we ensure that the reference is not influenced by the covariates through any of its ancestor nodes. If no suitable reference feature is known a priori, tascCODA provides an automatic way of selecting the feature with minimal dispersion across all samples among the features that are present in at least a share of samples *t* (default *t* = 0.95; this value can be lowered if no suitable feature exists).

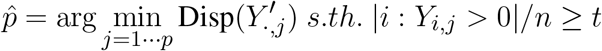

The restriction to large presence avoids choosing a rare feature as the reference where small changes in terms of counts lead to large relative deviations. The least-dispersion approach is aimed at reducing the bias introduced by the choice of reference. Equations (1–9) together with the reference feature yields the tascCODA model (Figure 1B):

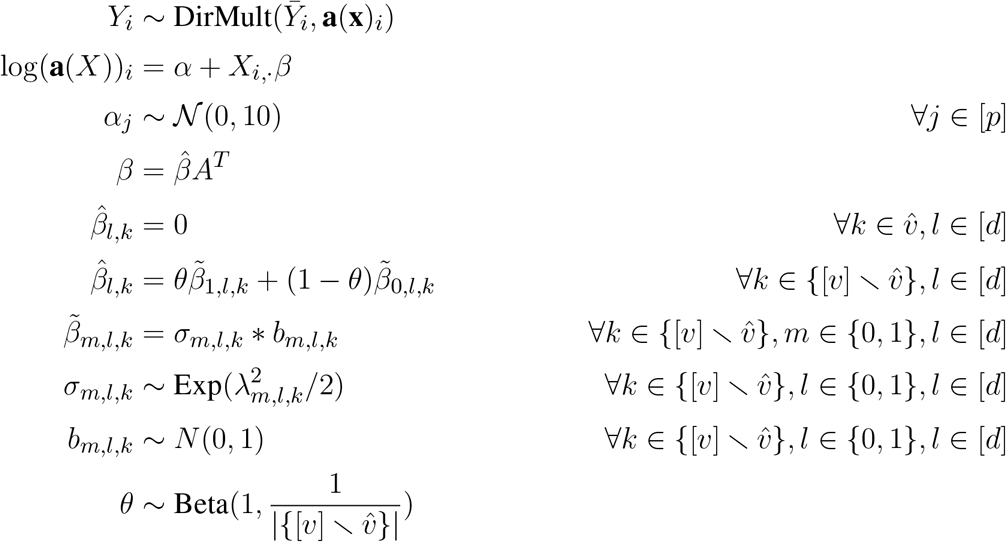

with the default choices of *λ*_0_ = 50 and *λ*_1,*k*_ set according to (10) with hyperparameters *φ* and *λ*_1_ = 5 (Supplementary material section 1.2).

### 2.2 Computational aspects

Before performing Bayesian inference with the tascCODA model, several data preprocessing steps are applied. Singular nodes, i.e., internal nodes that have only one child node, are removed from the tree, since their effect only propagates to one node and is therefore redundant. We also add a small pseudo-count of 0.5 to all zero entries of *Y* to minimize the frequency of numerical instabilities in our tests. Finally, we recommend normalizing all covariates to a common scale before applying tascCODA to avoid biasing the model selection process toward the covariate with the largest range of values.

Since tascCODA is a hierarchical Bayesian model, we use Hamiltonian Monte Carlo sampling (Betancourt and Girolami, 2015) for posterior inference, implemented through the tensorflow (Abadi et al., 2016) and tensorflow-probability (Dillon et al., 2017) libraries for Python, solving the gradient in each step via automatic differentiation. By default, tascCODA uses a leapfrog integrator with Dual-averaging step size adaptation (Nesterov, 2009) and 10 leapfrog steps per iteration, sampling a chain of 20,000 posterior realizations and discarding the first 5,000 iterations as burn-in, which was also the setting for all applications in this article, unless explicitly stated otherwise. As an alternative, No-U-turn sampling (Homan and Gelman, 2014) is available for use with tascCODA. The initial states for all *α_j_* and *b_m,l,k_* are randomly sampled from a standard normal distribution. All *σ_m,l,k_* and *θ* values are initialized at 1 and 0.5, respectively.

To determine the credible effects of covariates on nodes from the chain of posterior samples, we calculate the threshold of practical significance, introduced by Ročková and George (2018), for each node as follows:

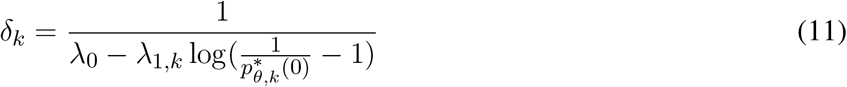

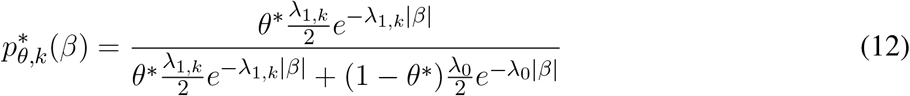

Here, *θ*^∗^ is the posterior median of *θ*. More details on *δ* are available in the Supplementary material. We compare the posterior median effects 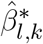 to the corresponding *δ_k_* and take all effects where 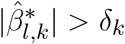 as credible. In the context of differential abundance testinf, we obtain the set of differentially abundant features *D* by multiplying the matrix with the all credible effects, 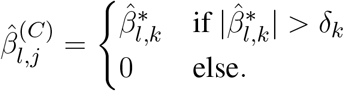, with *A^T^*, and get

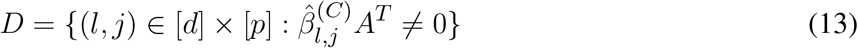

as the set of features, influenced by at least one credible effect.

A Python package for tascCODA is available at https://github.com/bio-datascience/tascCODA. Building upon the scCODA package, the software provides methods to seamlessly integrate scRNA-seq data from scanpy (Wolf et al., 2018) or microbial population data via pandas (McKinney, 2010). The package also allows to perform differential abundance testing with tascCODA and visualize tascCODA’s results through tree plots from the toytree package. All results were obtained using Python 3.8 with tensorflow=2.5.0 (Abadi et al. (2016)), tensorflow-probability=0.13 (Dillon et al. (2017)), arviz=0.11 (Kumar et al. (2019)), numpy=1.19.5, scanpy=1.8.1 (Wolf et al. (2018)), toytree=2.0.1, and sccoda=0.1.4 (Büttner et al. (2020)).

## 3 RESULTS

### 3.1 Simulation studies

#### 3.1.1 Model comparison

To test the performance of tascCODA in a differential abundance testing scenario, we generated compositional datasets with an underlying tree structure and compared how well several models could detect the changes introduced by a binary covariate. For compositional models that do not account for the tree structure, we used the state-of-the art methods ANCOM-BC (Lin and Peddada (2020)), ANCOM (Mandal et al. (2015)), and ALDEx2 (Fernandes et al. (2014)) from the field of microbiome data analysis, as well as scCODA (Büttner et al., 2020) from scRNA-seq analysis. Based on the recommendations by Aitchison (1982), we also analyzed the data with the additive log-ratio (ALR) transformation in combination with t- or Wilcoxon rank-sum tests. We also included the recent adaANCOM (Zhou et al., 2021a), a differential abundance testing method that accounts for the tree structure. Furthermore, we applied tascCODA with different values for the aggregation parameter, *φ* = (−10, −5, −1, 0, 1, 5, 10), setting *λ*_1_ = 5.

We first defined four different data sizes *p* = (10, 30, 50, 100) and randomly generated a multifurcating tree with depth 5 for each value of *p*. We then chose three nodes (one internal on the level directly above the leaves, two leaves) from each tree, whose child leaves, denoted by *p*′, are set to be differentially abundant under a binary (control-treatment) condition (Figure S1 - S4). Similar to Wadsworth et al. (2017), we generated *n* = *n*_0_ +*n*_1_ compositional data samples from two groups of equal size *n*_0_ = *n*_1_ = (5, 20, 30, 50). Each sample *Y_i_* is a realization of a Dirichlet-Multinomial distribution with a total sum of 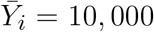 and a parameter vector *γ*^∗^. For extra dispersion in the data, we set 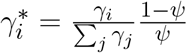 with *φ* = 0.002. The parameters for the first (control) group were generated via *γ*_0,*i*_ = exp (*α_i_*); *α_i_* ∼ Unif(2, 2). In the second (treatment) group, we added an effect *β* = (0.3, 0.5, 0.7, 0.9) to the components in 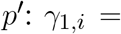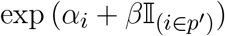. For each parameter combination (*p, n*_0_*, β*), we randomly generated 20 replicates, resulting in a total of 1280 datasets.

Since the adaANCOM method assumes a bifurcating tree structure, we transformed each tree node to a series of bifurcating splits via the *multi2di* and *collapse.singles* methods from the *ape* package for R (Paradis et al. (2004)) before applying the method. For the methods that require a reference category (ALR, scCODA, tascCODA, ALDEx2), we used the last component, which was always designed to be unaffected by the condition, as the reference. After applying each method to a dataset, we corrected the resulting p-values by the Benjamini-Hochberg procedure, except for ANCOM-BC, where we used the recommended Holm correction of p-values, and determined the significant results at an expected FDR level of 0.05. The Bayesian methods scCODA and tascCODA do not produce p-values and identify credible effects as previously described.

For an overall indicator of how well the different methods could determine differentially abundant features, we considered Matthews correlation coefficient (Figure 2A). Here, adaANCOM showed poor performance especially on small datasets, while ALDEx2 struggled when *p* was larger. Only scCODA and ANCOM-BC performed well in comparison for all data and effect sizes. For tascCODA, varying the aggregation level *φ* had a strong influence on the performance. With larger values of *φ*, tascCODA prefers less generalizing effects, resulting in a more detailed solution and larger MCC. At a high resolution level (*φ* = 5), tascCODA was on par with or even better than scCODA and ANCOM-BC, showing almost no sensitivity to the size of the dataset. Because the trees in our simulation contained only effects on leaf nodes or the level directly above, preferring generalizing effects (*φ* = −5) resulted in worse performance, while the unbiased case of *φ* = 0 gave slightly worse results than scCODA and ANCOM-BC. All methods shown in Figure 2B except adaANCOM controlled the FDR reasonably well, although ANCOM-BC and scCODA could not always hold the nominal level of 0.05. Only ALDEx2, which is known to be very conservative (Hawinkel et al., 2019; Büttner et al., 2020), produced almost no false positives, at the cost of larger type 2 error. tascCODA had a slightly inflated FDR (< 0.25) for smaller values of *φ* in some cases, which became more apparent when analyzing the ability of each method to exactly recover the true effects (2C). Increasing the effect size resulted in a reduced Hamming distance between the ground truth and tascCODA with *φ* = 5, which consistently outperformed all other models. tascCODA in the misspecified setting *φ* = −5 showed an inflated Hamming distance, especially for *p* = 30. This is, however, expected since tascCODA is forced to infer small-sized effects at the top level, resulting in many falsely detected features and thus a large deviation from the true sparse solution. In practice, this highlights the need to perform cross-validation over different levels of *φ* to reduce false discoveries due to misspecification. We further found that ANCOM detected many false positives in all of our simulations, while the ALR-based methods were similarly conservative as ALDEx2 (Figures **??**-**??**). Increasing the sample size generally improved the recovery performance of all methods except for tascCODA with misspecified *φ* (Figure **??**). 0.5cm

**Figure 2.**
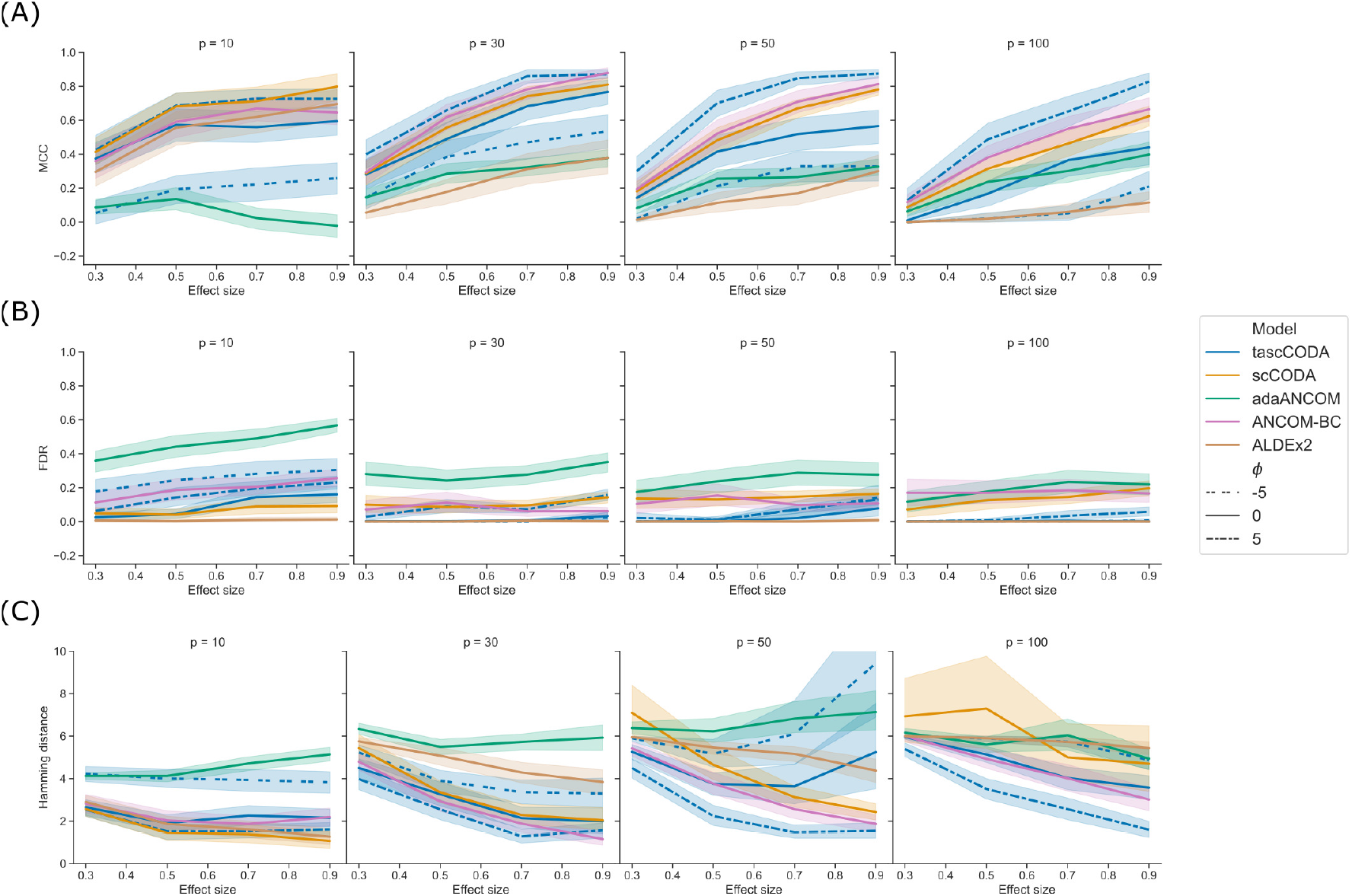
Performance comparison of tascCODA and other methods on simulated data with one binary covariate (differential abundance testing). Plots are grouped by the number of simulated components *p* and the effect size *β*. For tascCODA, different values of *φ* were tested (dashed blue lines). The areas around each line represent the standard deviation. Performance measured by **(A)** Matthews correlation coefficient (MCC). **(B)** False discovery rate (FDR) **(C)** Hamming distance between ground truth and determined effects.

#### 3.1.2 Effect detection at high tree levels

In the next benchmark scenario, we evaluated the effect of the tuning parameter *φ* in tascCODA to detect effects on larger groups of features through aggregation at higher levels of the tree. To this end, we considered the *p* = 30 setting with the tree structure from Figure S5, and defined an effect on a node near the root, influencing almost all features. We simulated datasets in the same manner as for the previous benchmark, with *n* = 10, *β* = (0.3, 0.5, 0.7, 0.9), and 20 replicates per effect size. We then compared tascCODA with different levels of *φ* using the same performance metrics as before.

With a correctly specified parametrization *φ* < 0, favoring effects near the root, tascCODA recovered almost all relevant effects, as indicated by a small Hamming distance and high MCC, without producing false positive results (Figure 3). With increasing *φ*, however, tascCODA favors effects on the leaves, thus entering the misspecified regime. As predicted, tascCODA was able to only recover a small portion of the true effects, while producing more false positive results. This highlights tascCODA’s ability to consistently uncover effects on larger groups of features which would be missed when not taking into account tree information.

**Figure 3.**
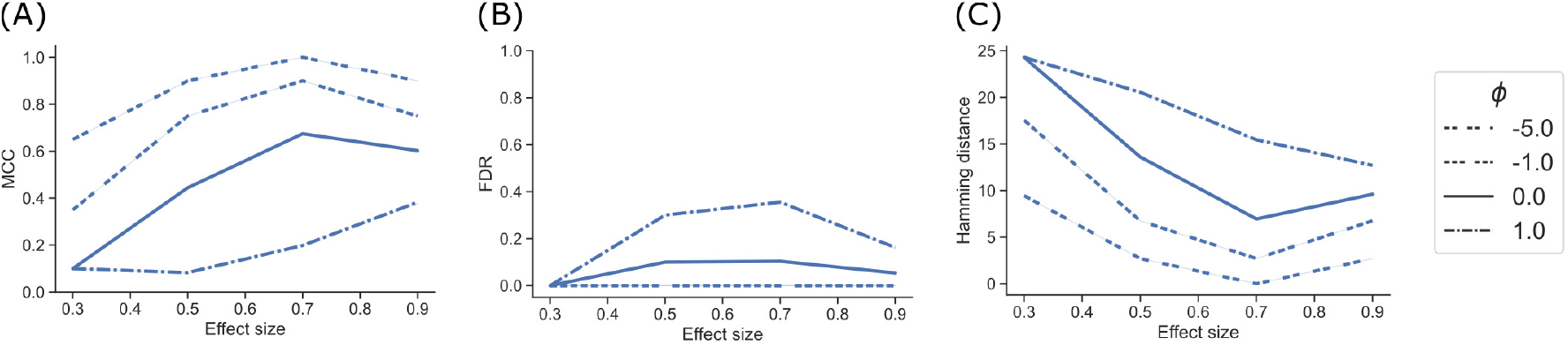
Performance comparison of different bias settings for tascCODA on simulated data with the effect being located near the root of the tree, depending on effect size. Performance measured by **(A)** Matthews correlation coefficient (MCC). **(B)** False discovery rate (FDR) **(C)** Hamming distance between ground truth and determined effects.

#### 3.1.3 Simulation with multiple covariates

In our third benchmark scenario, we simulated data with two covariates to showcase how tascCODA is able to distinguish effects from two different sources. Taking the tree from the method comparison study with *p* = 30 (Figure S2), we first defined a binary covariate *x*_0_ with effect sizes *β*_0_ = (0.3, 0.5, 0.7, 0.9) as before, and *n* = 10 samples per group. We also included a second covariate *x*_1_ ∼ *Unif* (0, 1) with effect size *β*_1_ = 3 that affects node 39 and therefore features 13-23 in all samples. For each effect size, we simulated 10 datasets and applied tascCODA with *φ* = (−5, 0, 5) and two different design matrices *X*. For the first design matrix, we used only *x*_0_, while the second design matrix contained both *x*_0_ and *x*_1_ as covariates. We compared how well both configurations could recover the effects introduced by *x*_0_ in terms of MCC, FDR, and Hamming distance to the ground truth.

Ignoring *x*_1_ in the model design resulted in an overall worse performance of tascCODA for all metrics, all effect sizes for *x*_0_, and all values of *φ* (Figure 4). In every case it proved beneficial to include the second covariate in the model, resulting in almost no false positive detections of changes caused by the first covariate. Further, the two-covariate model achieved an MCC and Hamming distance that were similar to our simulations where only one covariate acted on the data (Figure 2). This proves that tascCODA is able to reliably identify the influence of multiple covariates on the count data.

**Figure 4.**
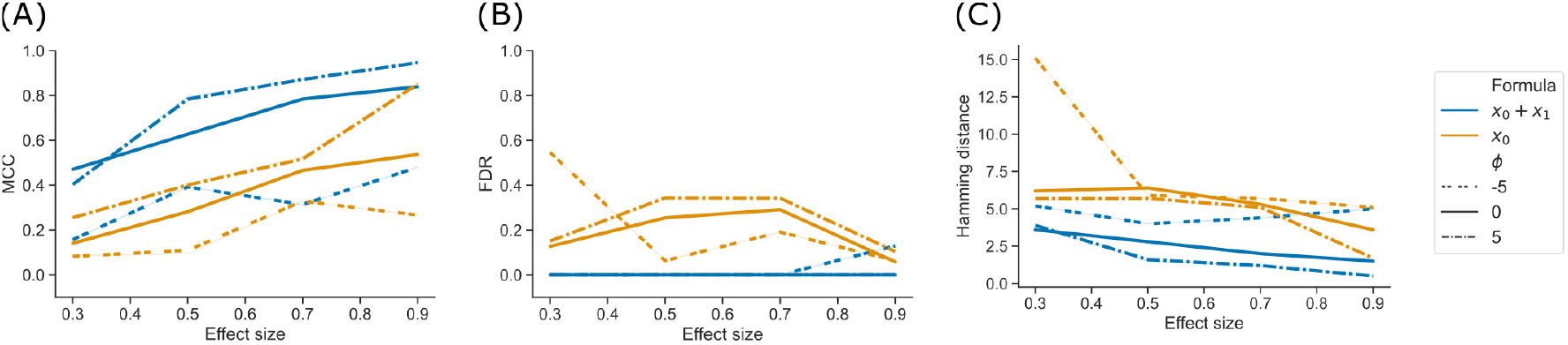
Performance comparison for tascCODA on simulated data with two covariates. The setups including both or only one covariate in the model are shown as *x*_0_ + *x*_1_ and *x*_0_, respectively. Simulations were evaluated for different effect sizes and aggregation levels *φ*. Performance measured by **(A)** Matthews correlation coefficient (MCC). **(B)** False discovery rate (FDR) **(C)** Hamming distance between ground truth and determined effects.

### 3.2 Experimental data applications

#### 3.2.1 Single-cell RNA-seq analysis of ulcerative colitis in humans

Ulcerative colitis is one of the most common manifestations of inflammatory bowel disease. The disease alternates between periods of symptomatic flares and remissions. The flares are due to the surge of an inflammatory reaction in the colon, causing superficial to profound ulcerations, which manifests with bloody stool, diarrhea and abdominal pain. The patients will thus have part of their colon referred to as “inflamed”, while colonic tissue still seemingly intact will be called “non-inflamed”. To show how tascCODA can be applied to cell population data from scRNA-seq experiments, we used data collected by Smillie et al. (2019) from a study of the colonic epithelium on ulcerative colitis (UC). In the study, a total of 133 samples from 12 healthy donors, as well as inflamed and non-inflamed tissue from 18 patients with UC, were obtained via single-cell RNA-sequencing, divided into epithelial samples and samples from the Lamina Propria (Supplemental data 1.3.1).

We applied tascCODA to six different subsets of the data, comparing two of the three health conditions in one type of tissue at a time, and then compared our findings with the results of scCODA and the Dirichlet regression model used by Smillie et al. (2019), implemented in the *DirichletReg* package for R (Maier (2014)). For tascCODA and scCODA, we used the automatically determined reference cell types, which are identical for both models in all cases, and applied scCODA with an FDR level of 0.05. In the Dirichlet regression model, we adjusted the p-values by the Benjamini-Hochberg procedure, and selected differentially abundant cell types at a level of 0.05.

The cell lineage tree inferred from Smillie et al. (2019) (Figure 5) is divided into epithelial, stromal and immune cells at the top level (Figure 5). While the biopsies from the Epithelium contain mostly epithelial cells, and samples from the Lamina Propria consist of cells mostly from the other two lineages, both groups also include considerable amounts of cells from the other major lineages. We first compared scCODA and Dirichlet regression, which both do not take the tree structure into account, to tascCODA with *φ* = 5 (Figure 6), thus preferring a detailed solution with effects mainly located on leaf nodes, which approaches the leaf-only solutions of the other two methods. In this setting, tascCODA, scCODA and Dirichlet regression all determined mostly epithelial cells to shift in abundance between pairwise comparisons of healthy, non-inflamed, and inflamed tissue samples from the intestinal Epithelium (Figure 6A), and most changes in the Lamina Propria to be among stromal and immune cells (Figure 6B). When propagating the node effects of tascCODA with *φ* = 5 to the leafs via Equation 13, the differentially abundant cell types determined by tascCODA, scCODA, and Dirichlet regression were largely identical (Figure 6).

**Figure 5.**
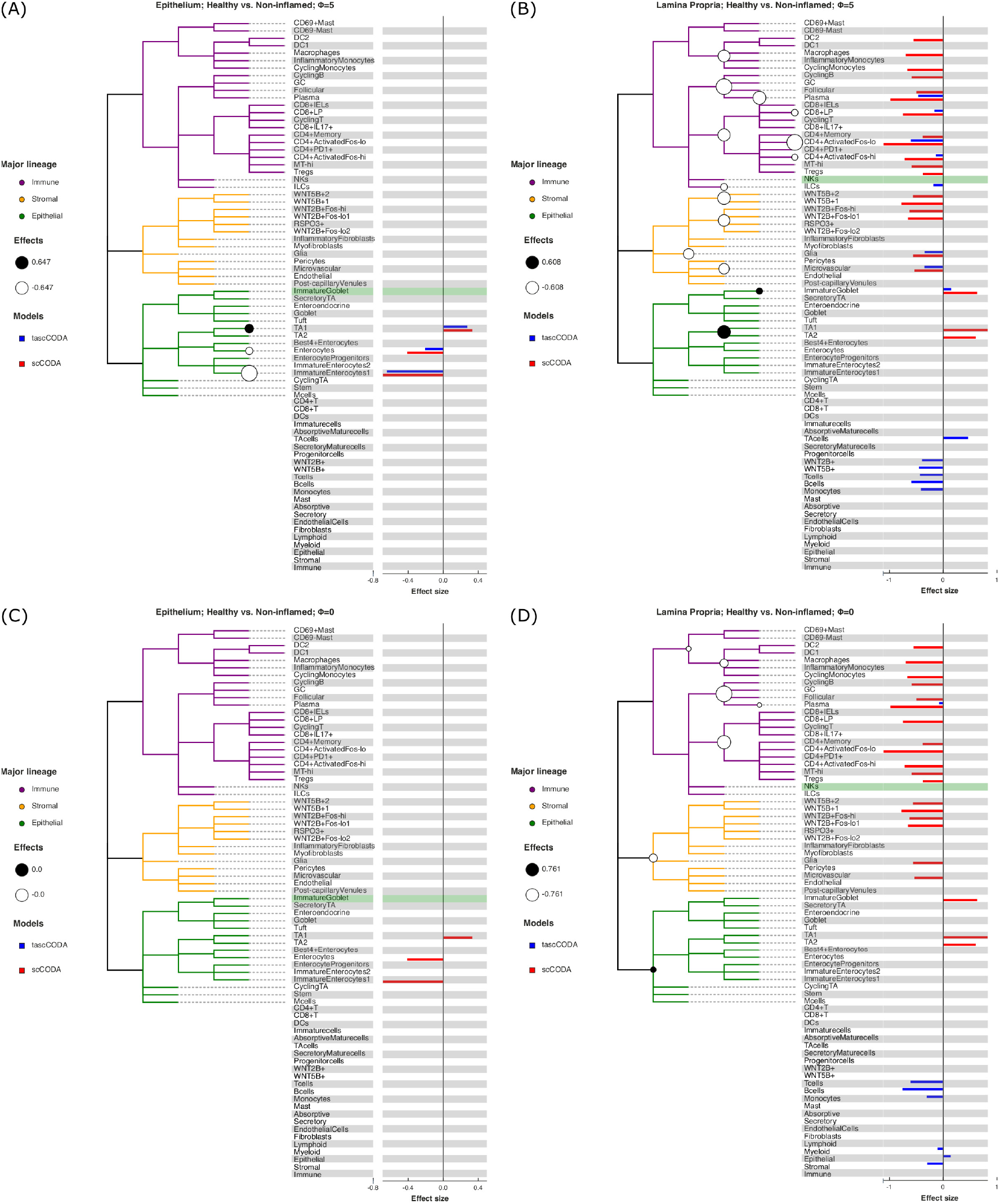
Behavior of tascCODA on scRNA-seq data for different values of *φ*. All plots show the comparison of healthy control samples to non-inflamed tissue samples of UC patients in the data from Smillie et al. (2019). White and black circles on the cell lineage tree show the effects found by tascCODA, which are also shown as blue bars on the right side of each plot. The bars below the tree depict effects on internal nodes, with lower positions in the diagram corresponding to nodes closer to the root. For comparison, the red bars indicate effects found by scCODA, which only operates on the tips of the tree. The green-shaded area shows the reference cell type that was used for both models. **(A)** When *φ* = 5, tascCODA prefers placing effects near the tips of the tree and finds the exact same solution as scCODA for the Epithelium data. **(B)** In the Lamina Propria, tascCODA places some effects on internal nodes, resulting in a sparser solution than the one obtained by scCODA (14 vs. 21 credible effects). **(C)** When *φ* = 0, tascCODA finds no credible effects in samples from the Epithelium, and **(D)** only seven effects are necessary to summarize the large number of effects found by scCODA when looking at samples from the Lamina Propria.

**Figure 6.**
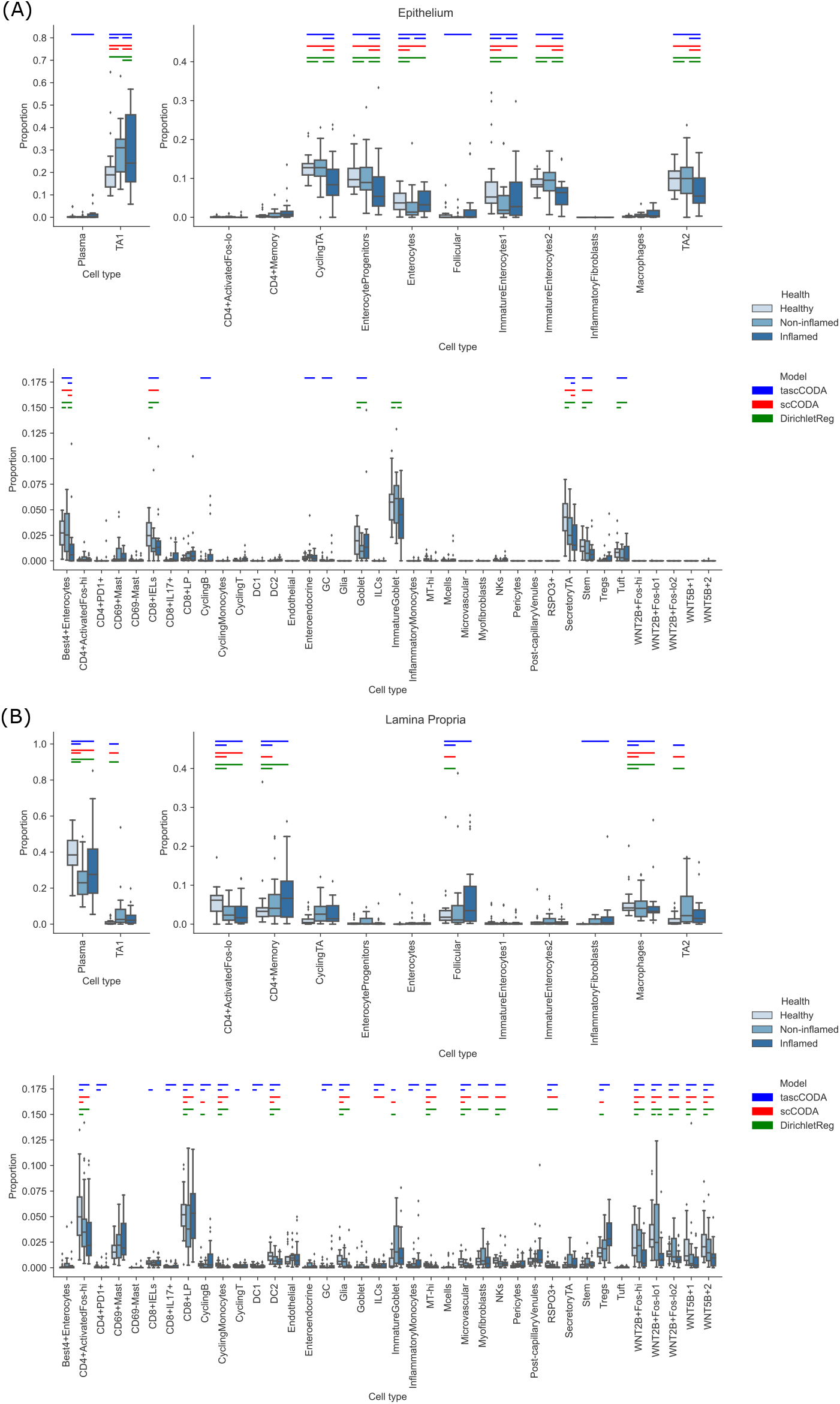
Comparison of differentially abundant cell types found by tascCODA (blue, *φ* = 5), scCODA (red, FDR=0.05), and Dirichlet regression (green, adjusted *p_adj_* < 0.05) between biopsies of healthy, non-inflamed and inflamed tissue. Colored bars for each method indicate that a credible change was found. **(A)** Among samples from the intestinal epithelium, tascCODA and Dirichlet regression detect effects on lowly abundant epithelial cell types (Tuft, Goblet, Enteroendocrine) that were not detected by scCODA. **(B)** In the Lamina Propria, only tascCODA detects a number of effects on some of the T and B cell types.

To further investigate the predictive and sparsity-inducing powers of tascCODA, we performed out-of-sample prediction with the results obtained from tascCODA and scCODA on 5-fold cross validation splits of each of the six data subsets. For both models, we determined cell type-specific effect vectors 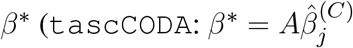, as in equation 13; scCODA: Model output) as well as the posterior mean of the base composition *α*^∗^ on the training splits, and used them to predict cell counts for each health status label *X_l_* in the corresponding test split as 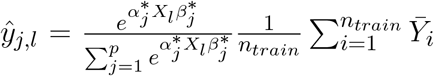. We measured the predictive power of tascCODA and scCODA as the mean squared logarithmic error (MSLE) between the actual and predicted cell counts, and sparsity as the average number of nonzero effects over all five splits (Table 1). For small *φ*, tascCODA determined very few or no credible effects, while the MSLE was usually slightly higher than the MSLE from scCODA. In the unbiased setting *φ* = 0, tascCODA found credible effects in three scenarios, which considerably reduced the MSLE. With a small bias towards the leaves (*φ* = 1), tascCODA even outperformed scCODA in terms of MSLE in one case, while for *φ* = 5, tascCODA achieved a lower MSLE and similar number of credible effects in three scenarios, and a lower number of credible effects and similar MSLE in the other three scenarios. We observed a curious result when comparing non-inflamed and inflamed epithelial samples. Here, the MSLE increased with rising *φ*, indicating that the mean model over all samples described the data better than trying to determine variation between the two groups. This confirms the intuition that the aggregation bias *φ* in tascCODA acts as a trade-off between generalization level and prediction accuracy. For smaller *φ*, tascCODA will select fewer, more general effects, which might miss subtle changes at a lower level of the lineage tree, while with increasing *φ*, tascCODA’s results will approach the ones discovered without taking tree aggregation into account.

**Table 1.** Mean squared logarithmic error (MSLE) and number of selected effects over 5 cross-validation splits for tascCODA with different parametrizations *φ* and scCODA. Abbreviations for scenarios: Healthy (H), Non-inflamed (N), and Inflamed (I). With increasing *φ*, tascCODA selects more effects and on average improves its predictive power. At *φ* = 5, tascCODA has equal or lower MSLE than scCODA and a similar number of selected effects

| Scenario | Model<br>$\phi$ | tascCODA | | | | | scCODA |
| --- | --- | --- | --- | --- | --- | --- | --- |
|  |  | -5 | -1 | 0 | 1 | 5 | - |
| Epithelium - H vs. N | MSLE | 142.22 | 142.16 | 142.18 | 138.56 | 134.36 | 134.96 |
|  | Effects | 0.0 | 0.0 | 0.0 | 1.2 | 3.2 | 2.4 |
| Epithelium - H vs. I | MSLE | 167.46 | 163.60 | 160.68 | 158.06 | 154.64 | 154.44 |
|  | Effects | 0.0 | 1.6 | 2.6 | 3.2 | 8.2 | 10.8 |
| Epithelium - N vs. I | MSLE | 173.94 | 174.10 | 174.10 | 175.86 | 177.26 | 174.78 |
|  | Effects | 0.0 | 0.0 | 0.0 | 0.2 | 3.6 | 5.2 |
| LP - H vs. N | MSLE | 162.76 | 157.62 | 155.16 | 152.80 | 149.58 | 154.02 |
|  | Effects | 0.4 | 1.8 | 3.0 | 6.2 | 16.0 | 14.4 |
| LP - H vs. I | MSLE | 188.58 | 182.96 | 178.88 | 176.02 | 173.32 | 173.40 |
|  | Effects | 0.0 | 1.8 | 4.8 | 7.8 | 17.8 | 17.4 |
| LP - N vs. I | MSLE | 219.72 | 219.70 | 219.66 | 219.68 | 216.76 | 218.62 |
|  | Effects | 0.0 | 0.0 | 0.0 | 0.0 | 1.4 | 0.4 |

For a more detailed comparison between tascCODA and scCODA, we compared healthy to non-inflamed biopsies of control and UC patients. When choosing *φ* = 5, thus biasing tascCODA towards the leaf nodes, tascCODA detected the differences in cell composition in the Epithelium as changes in abundance of the same three cell types as scCODA (Figure 5A). In the Lamina Propria, tascCODA detected credible changes on six different groups of cell types, including T and B cells, which were previously linked to UC (Holmén et al. (2006); Smillie et al. (2019)), as well as eight single cell types (Figure 5B). Notably, tascCODA amplified the decrease of Plasma B-cells induced by the group effect on B-cells by an additional negative effect on the cell type level. A strong decrease of Plasma cells was also confirmed by Smillie et al. (2019) through FACS stainings. Importantly, tascCODA described the data with only 14 nonzero effects, whereas with scCODA, 21 credible effects were produced.

As a contrast, we also examined the unbiased setting with *φ* = 0, treating all nodes equally. Here, the cell type-specific changes in the Epithelium were not picked up anymore by tascCODA (Figure 5C). In the Lamina Propria, only seven effects, almost all on groups of cell types, were detected by tascCODA (Figure 5D). Again, B and T cells were found as the cell lineages that undergo the largest change between healthy and non-inflamed UC biopsies. When testing healthy versus inflamed, and non-inflamed versus inflamed biopsies, tascCODA also detected more detailed results when *φ* = 5, and found fewer, more generalizing effects with *φ* = 0 (Figure **??**, **??**; Table **??**-**??**).

#### 3.2.2 Analysis of the human gut microbiome under Irritable Bowel Syndrome

We next considered a microbiome data example and considered another chronic disorder of the human gut, the Irritable Bowel Syndrome (IBS). IBS is a functional bowel disorder characterized by frequent abdominal pain, alteration of stool morphology and/or frequency, with the absence of other gastrointestinal diseases (i.e. colorectal cancer, inflammatory bowel disease). It is estimated that about 10% of the general population experience symptoms that can be classified as a subtype of Irritable Bowel Syndrome, which include IBS-C (constipation), IBS-D (diarrhea), IBS-M (mixed), or unspecified IBS (Ford et al. (2017)). While the exact sources of the disease can be manifold, it has been hypothesized that the gastroenterological symptoms may be caused by a disturbed composition of the gut microbiome (Duan et al. (2019); Ford et al. (2017)).

In particular, we analyzed 16S rRNA sequencing data of stool samples collected from IBS patients and healthy controls, which were obtained by Labus et al. (2017). The dataset consists of *n* = 52 samples, with 23 healthy controls, and 29 IBS patients separated into 11 subjects with constipation (IBS-C), 10 subjects with diarrhea (IBS-D), 6 subjects with mixed symptoms (IBS-M), and 2 subjects with unspecified symptoms. Further, metadata information about age, sex and BMI of most subjects is available. We re-processed the raw 16S rRNA sequences with DADA2, version 1.21.0 (Callahan et al. (2016)) and did taxonomic assignment via the Silva database, version 138.1 (Quast et al. (2013); Yilmaz et al. (2014)), yielding a final count table with 709 ASVs along with a taxonomic tree (Supplemental data 1.3.2). This data was then aggregated at the genus level, resulting in a total of *p* = 91 known genera.

We applied tascCODA to the genus-level data, comparing healthy and IBS subjects. For comparison, we also applied scCODA and ANCOM to the data aggregated at each level of the taxonomic tree (phylum, class, order, family, and genus). To showcase the flexibility of tascCODA, we analyzed the data with different covariate setups, by including the other available metadata variables. As a reference genus for scCODA and tascCODA, we chose *Alistipes*, since it is a genus with relatively high presence and rather low dispersion. For all analyses on this dataset, we decreased the mean shrinkage in tascCODA to *λ*_1_ = 1, allowing us to find more subtle effects.

We first used tascCODA to analyze the differences in the gut microbial composition between healthy controls and IBS patients (Figure 7, Table **??**). Favoring generalization with *φ* = −5, we found only a small decrease of the phylum Firmicutes (Figure 7A). In the unbiased setting (*φ* = 0), the previous effect on the phylum level was substantiated to the Oscillospirales order. Additionally, decreases of the *Parabacteroides* and *Bacteroides* genera are found (Figure 7B). Setting *φ* = 5, thus favoring detailed results, we discovered a decrease of the Ruminococcaceae family, a subgroup of Oscillospirales, and multiple decreasing genera with the strongest effects on *Parabacteroides* and *Bacteroides* (Figure 7C). For comparison, we also applied scCODA (FDR=0.1) to the same dataset, which also discovered a decrease of *Parabacteroides* and *Bacteroides*, as well as three genera in the Ruminococcaceae family. A decrease of *Parabacteroides* in a subset of IBS patients was also found by Labus et al. (2017). Also, a relative decrease of the order Bacteroidales, which includes *Parabacteroides* and *Bacteroides*, was reported by Nagel et al. (2016) and Jeffery et al. (2012). Decreasing shares of Ruminococcaceae were also connected to IBS in multiple studies (Pozuelo et al., 2015; Durbán et al., 2012).

**Figure 7.**
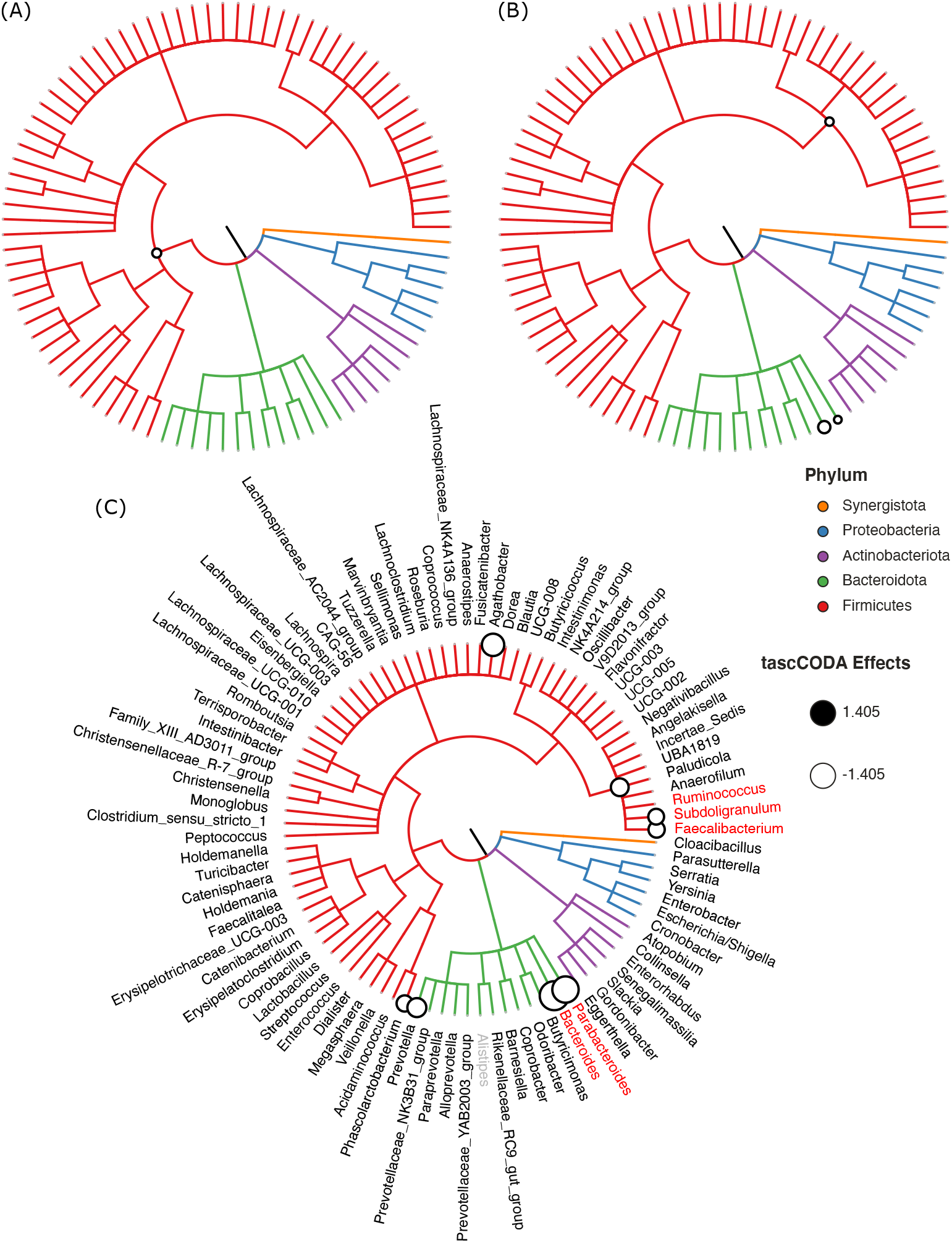
Credible changes found by tascCODA (*λ*_1_ = 1), comparing healthy controls and IBS patients in the genus-aggregated data of Labus et al. (2017). The circles on nodes of the tree represent credible effects. **(A)** High-level aggregation with *φ* = 5. **(B)** Unbiased aggregation (*φ* = 0). **(C)** Aggregation with bias towards the leaves (*φ* = 5). Red genera show the credible effects found by scCODA (FDR=0.1) on the genus level. The grey genus *Alistipes* was used as the reference for tascCODA and scCODA.

To highlight the flexibilty of tascCODA, we next tried to discover changes in the gut microbiome related to age, BMI, gender, and IBS subtype. Before applying tascCODA, we min-max normalized the two former covariates to obtain a common scale for all covariates. We excluded three samples with missing information on BMI. We conducted every analysis three times with *φ* = −5, 0, 5. When testing for changes related to one of age, gender, or BMI alone, tascCODA was not able to discover any credible differences for any aggregation bias. When testing on all four covariates together, excluding interactions, tascCODA only reported credible changes in the microbiome with respect to the IBS subtype. Finally, including all possible variable, interactions revealed that while a general negative effect was found independent of gender, male IBS-D patients had a larger depletion of *Bacteroides* than female patients.

Next, we restricted our analysis to testing for changes between the four IBS subtypes and all other samples. The results shown in Figure 8 and Table **??** were obtained with *φ* = 5. For patients experiencing constipation (IBS-C, Figure 8A), decreases of *Agathobacter*, *Bacteroides*, *Ruminococcus*, and *Faecalibacterium*, as well as an increase of *Anaerostipes* were found by tascCODA. Conversely, diarrhea (IBS-D, Figure 8B) was associated with a decrease in *Parabacteroides*, as well as a large decrease in *Bacteroides*. Patients with mixed symptoms (IBS-M, Figure 8C) were found to have increased numbers of *Blautia*, in addition to a decrease of *Parabacteroides* and *Faecalibacterium*, which each match with the observations related to one of the two previous conditions. Finally, only a small increase of *Romboutsia* was associated to IBS with unspecified symptoms (IBS-unspecified, Figure 8D).

**Figure 8.**
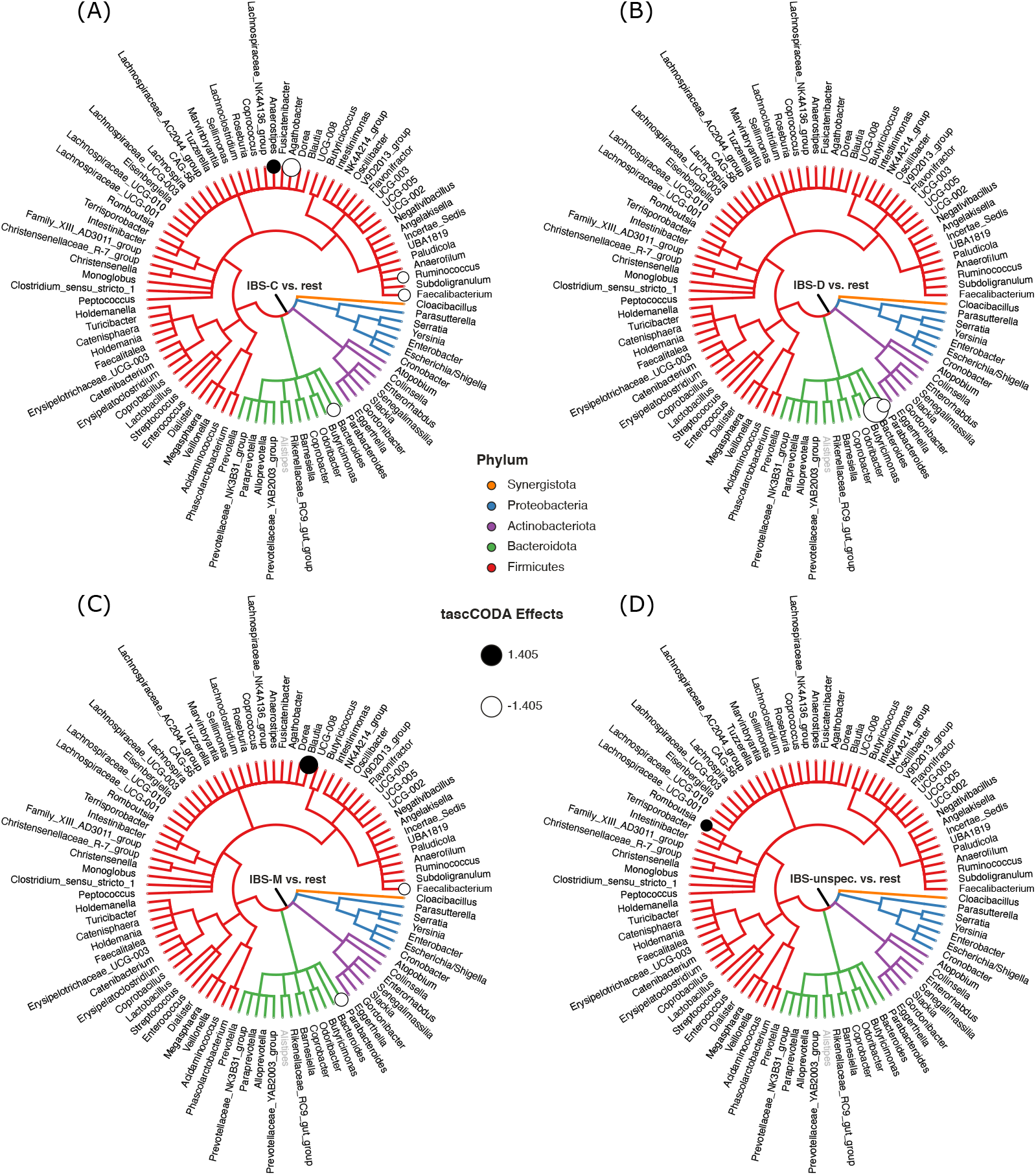
Credible changes found by tascCODA (*λ*_1_ = 1*, φ* = 5), simultaneously comparing healthy controls to all IBS subtypes in the genus-aggregated data of Labus et al. (2017). The circles on nodes of the tree represent credible effects. The grey genus *Alistipes* was used as the reference for tascCODA. **(A)** IBS-C (n=11). **(B)** IBS-D (n=10). **(C)** IBS-M (n=6). **(D)** IBS-unspecified (n=2).

## 4 DISCUSSION

Associating changes in the structure of microbial communities or cell type compositions with host or environmental covariates are commonly investigated with amplicon or single-cell RNA sequencing. With tascCODA, we have presented a fully Bayesian method to determine such compositional changes that acknowledges the hierarchical structure of the underlying microbial or cell type abundances and simultaneously accounts for the compositional nature of the data. By introducing tree-based penalization that adapts to the structure of the tree, the tascCODA model is able to accurately identify group-level changes with fewer parameters than traditional individual feature-based approaches. Thanks to a scaled variant of the spike-and-slab lasso prior (Ročková and George (2018)), we were able to obtain sparse solutions that can favor high-level aggregations or more detailed effects on a dynamic range characterized by a single scaling parameter *φ*. The tascCODA Python package seamlessly integrates into the *scanpy* environment for scRNA-seq (Wolf et al. (2018)) and allows Bayesian regression-like analyses with flexible covariate structures.

Through its ability to favor general trends or more detailed solutions, tascCODA is able to provide a trade-off between model sparsity and accuracy, which can be adjusted to reveal credible associations on different levels of the hierarchy. We recapitulated this behavior in synthetic benchmark scenarios, where focusing on low aggregation levels allowed tascCODA to outperform state-of-the-art methods in a differential abundance testing setup, while effects that influenced the majority of features were recovered with greater accuracy when we favored generalizing solutions. The aggregation property further allows for more interpretable models, detecting group-specific changes in the cell lineage or microbial taxonomy. For instance, tascCODA determined B and T cells as the main factors in cell composition changes of the Lamina Propria of Ulcerative Colitis patients, while inflamed epithelial tissue biopsies showed a depletion of Enterocytes.

Second, tascCODA can accommodate any linear combination of normalized covariates, allowing for multi-faceted analysis of complex relationships, while still producing highly sparse and interpretable solutions. On synthetic data, we showed that tascCODA was able to accurately distinguish the influence of two covariates that perturbed the data in different ways. While we did not detect credible relationships with the covariates age, sex and BMI, tascCODA was also able to simultaneously identify characteristic shifts in the gut microbiome for each subtype of Irritable Bowel Syndrome.

The application range of tascCODA extends beyond the taxonomic or expert-derived cell lineage tree structures used in our real data applications. Genetically driven orderings such as phylogenetic trees or cell type hierarchies obtained from clustering algorithms, or fully correlation-based approaches may provide more accurate results in differential abundance testing (see, e.g., Bichat et al. (2020) for further information).

While tascCODA provides a hierarchically adaptive extension of a classical compositional modeling framework based on a fixed aggregation level, extensions of the method could increase the application range of tascCODA. First, tascCODA does not account for the zero-inflation and overdispersion that is common in microbial abundance data on the OTU/ASV level. We avoided this challenge here by aggregating to the genus level. Accounting for these properties within the model, for example by using a zero-inflated Dirichlet-Multinomial model (Tang and Chen (2019)) or the Tweedie family of distributions (Mallick et al. (2021)), would allow for even more fine-grained analyses. Second, the tascCODA model currently places a sparsity-inducing spike-and-slab lasso prior on all included covariates. A natural next step would be to consider some covariates as confounding variables similar to Zhou et al. (2021b), reducing the number of latent parameters, while restricting results to a few core influence factors. Third, extending known efficient computational methods for inference of spike-and-slab lasso priors (Bai et al. (2020b); Ročková and George (2018)) to be used with our compositional modeling framework could greatly reduce the computational resources required for running tascCODA.

We believe that tascCODA, together with its implementation in Python, represents a valuable addition to the growing toolbox of compositional data modeling tools by providing a unifying statistical way to model and analyze microbial and cell population data in the presence of hierarchical side information.

## AUTHOR CONTRIBUTIONS

JO developed tascCODA and conducted the simulation studies and real data analysis. SC processed the 16S rRNA sequencing data and provided biological context. CLM supervised the work. JO and CLM conceived the statistical model, designed the simulation and out-of-sample prediction studies and wrote the manuscript. All authors read and approved the final manuscript.

## ACKNOWLEDGMENTS

We thank Dr. Maren Büttner for providing the initial processing steps in the scRNA-seq data analysis. Furthermore, we thank Dr. Jennifer S. Labus for kindly sharing additional metadata information on the IBS data. We acknowledge Dr. Michael Menden’s support in supervising SC during her Master’s Thesis.

## SUPPLEMENTAL DATA

Supplementary Material should be uploaded separately on submission, if there are Supplementary Figures, please include the caption in the same file as the figure. LaTeX Supplementary Material templates can be found in the Frontiers LaTeX folder.

## DATA AVAILABILITY STATEMENT

The model is available as a Python package on github^4^. The datasets used in this study are publicly available on Single Cell Portal (accession ID SCP259) and the Short Read Archive (accession number PRJNA373876). The scripts used for data analysis and benchmark data generation can be found in the tascCODA reproducibility repository^5^. Supplemental data can be downloaded from zenodo^6^.

## Footnotes

1 available at https://github.com/bio-datascience/tascCODA

2 https://github.com/bio-datascience/tascCODA

3 https://zenodo.org/record/5302136#.YSrhdi1h0mI

4 https://github.com/bio-datascience/tascCODA

5 https://github.com/bio-datascience/tascCODA_reproducibility

6 10.5281/zenodo.5302135

