## Supplementary Material for "tascCODA: Bayesian tree-aggregated analysis of compositional amplicon and single-cell data"

### 1 SUPPLEMENTARY DATA

#### 1.1 Notation overview

This section gives an overview over the inputs and parameters used by tascCODA:

**Data inputs**

- $Y \in \mathbb{R}^{n \times p}$ is the count matrix of features $j = 1 \dots p$ in samples $i = 1 \dots n$. $\bar{Y}_i = \sum_{j=1}^{p} Y_{i,j}$ is the sequencing depth of sample $i$.
- $X \in \mathbb{R}^{n \times d}$ is the covariate matrix of covariates $l = 1 \dots d$ for samples $i = 1 \dots n$.
- $\mathcal{T}$ is a multifurcating tree structure with $p$ leaves and $t$ internal nodes defined by the ancestor matrix $A \in \{0, 1\}^{p \times v}$, with $v = p + t$

**Latent parameters**

- $\mathbf{a}_i = (a_{1,i}, \dots a_{p,i}); a_{j,i} \geq 0$ is the probability vector of the Dirichlet-Multinomial distribution for sample $i$.
- $\alpha_j$ is the base (intercept) parameter for feature $j$.
- $\beta_{l,j}$ is the effect of covariate $l$ on feature $j$.
- $\hat{\beta}_{l,k}$ is the effect of covariate $l$ on tree node $k$.
- $\tilde{\beta}_{0,l,k}$ is the spike portion of the spike-and-slab LASSO prior for covariate $l$ and tree node $k$ with parameters $\sigma_{0,l,k}$ and $b_{0,l,k}$.
- $\tilde{\beta}_{1,l,k}$ is the slab portion of the spike-and-slab LASSO prior for covariate $l$ and tree node $k$with parameters $\sigma_{1,l,k}$ and $b_{1,l,k}$.
- $\theta$ is the mixture coefficient of the spike-and-slab LASSO prior.

**Tuning parameters/hyperparameters**

- $\lambda_0$ is the shrinkage parameter for the spike portion, default $\lambda_0 = 50$.
- $\lambda_{1,k}$ is the node-specific shrinkage parameter for the slab portion of the prior on node $k$, with mean value $\lambda_1$, default $\lambda_1 = 5$.
- $\phi$ is the aggregation bias parameter for scaling the slab shrinkage $\lambda_{1,k}$

#### 1.2 Hyperparameters for the spike-and-slab LASSO prior

We want to shed some additional light on the role of the hyperparameters $\lambda_0, \lambda_1, \theta$ in the spike-and-slab LASSO prior (Ročková and George (2018)). For simplicity and because the model is symmetric with respect to the covariates, we assume $d = 1$ and thus refrain from indexing parameters with the covariate. For one node $\hat{\beta}_k$, the prior is a mixture of two double-exponential distributions $\psi_0(\hat{\beta}_k)$ and $\psi_1(\hat{\beta}_k)$ (Figure S1A) whose share is determined by $\theta$:

$$p(\hat{\beta}_k|\theta) = \theta\psi_1(\hat{\beta}_k) + (1-\theta)\psi_0(\hat{\beta}_k) \tag{S1}$$

$$\psi_1(\hat{\beta}_k) = \frac{\lambda_1}{2}e^{-\lambda_1|\hat{\beta}_k|} \tag{S2}$$

$$\psi_0(\hat{\beta}_k) = \frac{\lambda_0}{2}e^{-\lambda_0|\hat{\beta}_k|} \tag{S3}$$

The double-exponential density (S2) has a peak at zero for large values of $\lambda$, which decreases with $\lambda$ (Bai et al. (2020)). Thus, setting $\lambda_0 \gg \lambda_1$ in the mixture density (S1) results in a product of a peaked "spike" ($\psi_0$) and a diffuse "slab" ($\psi_1$) component (Figure S1B). Interestingly, Ročková and George (2018) showed that the spike-and-slab LASSO prior can be reformulated as a penalized likelihood method that is, for fixed $\theta$:

$$pen(\hat{\beta}_k|\theta) = -\lambda_1|\hat{\beta}_k| + \log(\frac{p^*_\theta(0)}{p^*_\theta(\hat{\beta}_k)}) \tag{S4}$$

where

$$p^*_\theta(b) = \frac{\theta\frac{\lambda_1}{2}e^{-\lambda_1|b|}}{\theta\frac{\lambda_1}{2}e^{-\lambda_1|b|} + (1-\theta)\frac{\lambda_0}{2}e^{-\lambda_0|b|}} \tag{S5}$$

In the case of $\lambda_0 = \lambda_1$, the log-term in (S4) vanishes, and the penalty is equivalent to the standard LASSO (Tibshirani (1996)).

After making the weight $\theta$ data-adaptive by a Beta prior (Equation (9)), we turn our attention to the double-exponential parameters. We show the influence of each parameter on the solution by simulations on one of the randomly generated datasets from the simulation study with $p = 10$ features. From Figure S2, we can see that the ground truth assumption are effects on nodes 0, 4, and 12, with the latter node affecting features 7 and 8. We first fix $\lambda_1 = 1$, and vary $\lambda_0$ on a scale between 1 and 1000. Figure S1C shows that the effects $\hat{\beta}$ quickly stabilize with the three true effects being clearly separated from all other effects, which are close to zero. This stabilization was also explained by Ročková and George (2018) and is rooted in the fact that larger values of $\lambda_0$ only narrow the spike, which does not affect the solution after some point. We can thus simply set $\lambda_0$ to a relatively large value, the default in tascCODA is $\lambda_0 = 50$. When $\lambda_0 = \lambda_1$, we can see the typical parameter curve of a LASSO model, where the true effects are the last to approach zero (Figure S1D).

Because $\lambda_1 \to \lambda_0$ approaches the $\mathcal{L}_1$ penalty of the LASSO, which will eventually force all effects towards zero, leaving $\lambda_0 = 50$ and increasing $\lambda_1$ shows a similar behavior (Figure S1E). Only the true effects are significantly larger than zero once $\lambda_1$ reaches a value of approximately $0.1$. After a certain point ($\lambda_1 \approx 10$), the penalty becomes so large that all effects vanish. We utilize the regularizing behavior by scaling $\lambda_1$ depending on the number of leaves that a node influences to put a preference on nodes on different levels of the tree (Equation 10). The direction and steepness of the preference is expressed by the parameter $\phi$, with $\phi = 0$ giving equal treatment to all nodes. The default overall size of the penalty, $\lambda_1 = 5$, is chosen in a way that the parameters $\lambda_{1,k} \in (0, 10]$ stay in the range of values that were recommended by Ročková and George (2018) for all $k$. Figure S1F shows how the results change with different values of $\phi$.

For $\phi <= 0$, favoring high-level aggregations, the model selects the three ground truth nodes. When $\phi > 1$, tips are penalized considerably less than internal nodes and the effect on node 12 is replaced by equal-sized effects on its children, nodes 7 and 8. Also, for $\phi < 0$, effects on nodes that are high in the tree (large $k$) are different from zero, but smaller than the significance threshold (dashed line), while for $\phi > 0$, this is the case for leaf nodes.

### 1.3 Experimental data preprocessing

#### 1.3.1 Single-cell RNA-seq analysis of ulcerative colitis in humans

We obtained the data on ulcerative colitis from from Single Cell Portal (accession ID SCP259) and the analysis code from github. In total, the data consists of 365,492 transcriptomes from 12 healthy donors and 18 donors with UC providing non-inflamed and inflamed tissue samples. We used the 51 different cell types found in the original analysis, but considered every replicate as an independent sample, as done in a re-analysis by Büttner et al. (2020) on the same dataset. Biopsies from two different tissue regions, the Epithelium ('Epi' - 24 healthy, 21 non-inflamed, 16 inflamed samples) and the underlying Lamina Propria ('LP' - 24 healthy, non-inflamed, and inflamed samples each), were divided by enzymatic digestion. We inferred the cell lineage tree from the Methods section of Smillie et al. (2019) (Figure **??**).

#### 1.3.2 Analysis of the human gut microbiome under Irritable Bowel Syndrome

The raw 16S rRNA sequences (avilable at the Short Read Archive, accession number PRJNA373876) were re-processed using DADA2, version 1.21.0 (Callahan et al. (2016)). After primer and quality filtering (minimum read length: 150bp, maximum errors per read: 3, reads trimmed at first base with quality below: 10), inference of ASVs and removal of chimeras, the taxonomy of the inferred ASVs was determined with the Silva database, version 138.1 (Quast et al. (2013); Yilmaz et al. (2014)). Samples with a total read count of less than 500 (n=0) were discarded and ASVs assigned to Eukaryota (n=0) or belonging to an unknown Phylum (n=1) were removed, yielding a final count table with 709 ASVs along with a taxonomic tree.

### 2 SUPPLEMENTARY TABLES AND FIGURES

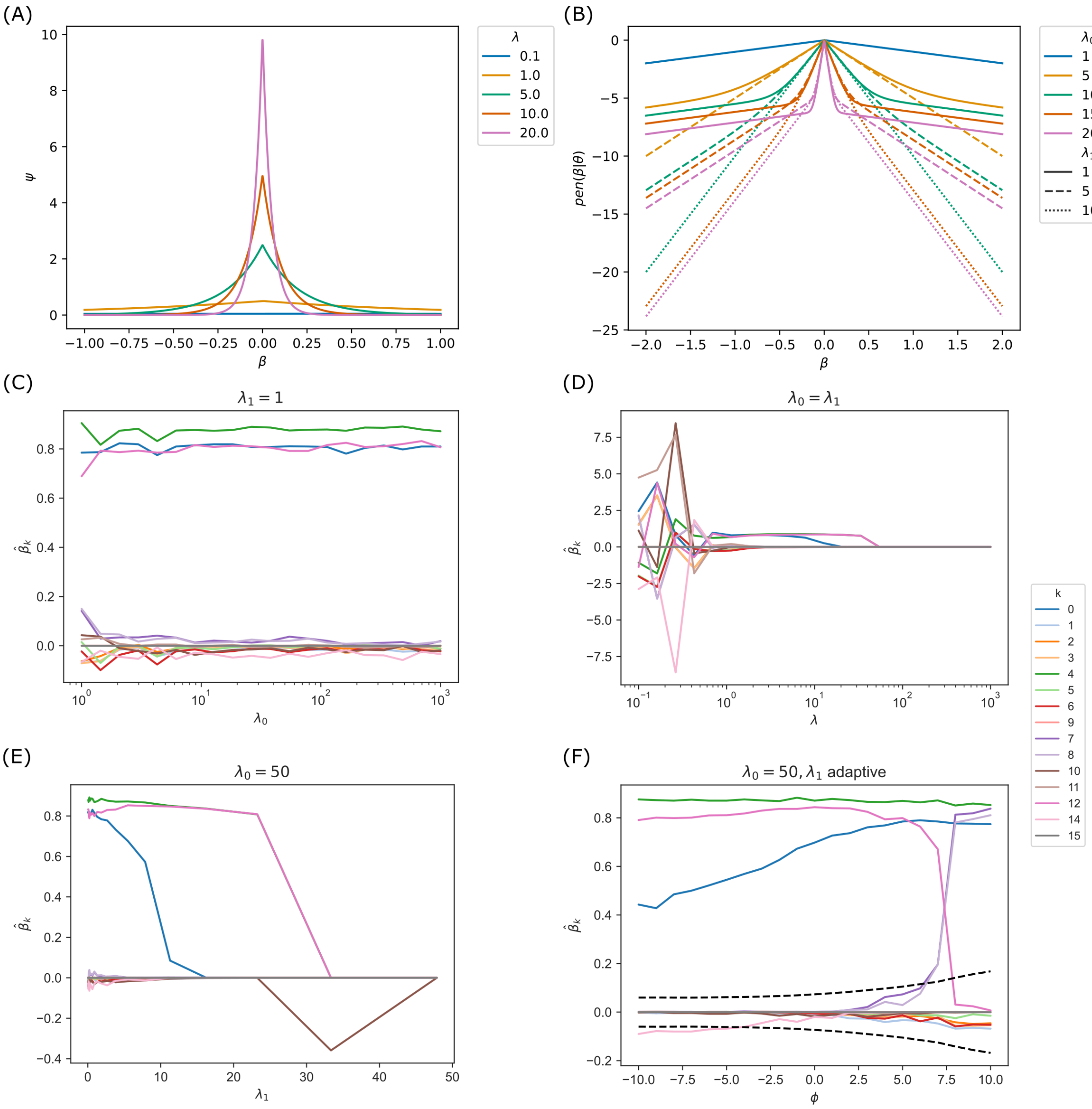


**Figure S1.** Parameters in the spike-and-slab LASSO penalty. **(A)** The double exponential density $\psi(\beta, \lambda)$ for different values of $\lambda$. The density becomes more peaked with increasing $\lambda$. **(B)** The likelihood penalty (Equation (S4)) introduced by different parametrizations of the spike-and-slab LASSO prior ($\theta = 0.1$). For larger effect sizes $\beta$, the penalty is driven by the slab parameter $\lambda_1$ (lines with the same style are close together). For smaller effect sizes $\beta$, the penalty is driven by the spike parameter $\lambda_0$ (lines with the same color are close together). If $\lambda_0 = \lambda_1$, the penalty is linear and equivalent to the LASSO penalty $\lambda_0\beta$. **(C-F)** Effect of different parameters on the effects $\hat{\beta}_k$ determined by tascCODA. For all simulations, a realization of the dataset in Supplementary Figure S2 was used. The nodes 13, 16 and 17 are singularities and were thus deleted before model application. **(C)** Solutions found by tascCODA when varying values of $\lambda_0$ and constant $\lambda_1 = 1$. The effects $\hat{\beta}_k$ stabilize and increasing $\lambda_0$ has no effect. **(D)** Solutions found by tascCODA in a LASSO-equivalent setting when varying values of $\lambda_0 = \lambda_1 = \lambda$. With increasing $\lambda$, more effects $\hat{\beta}_k$ go to 0. **(E)** Solutions found by tascCODA when varying values of $\lambda_1$ and constant $\lambda_0 = 50$. With increasing $\lambda_1$, a similar effect to the LASSO can be seen, where all effects are eventually approaching 0. **(F)** Solutions found by tascCODA when varying the tree level bias $\phi$. $\lambda_0 = 50, \lambda_{1,k}$ as in Equation 10. The dashed black lines show the significance threshold (Equation 11).

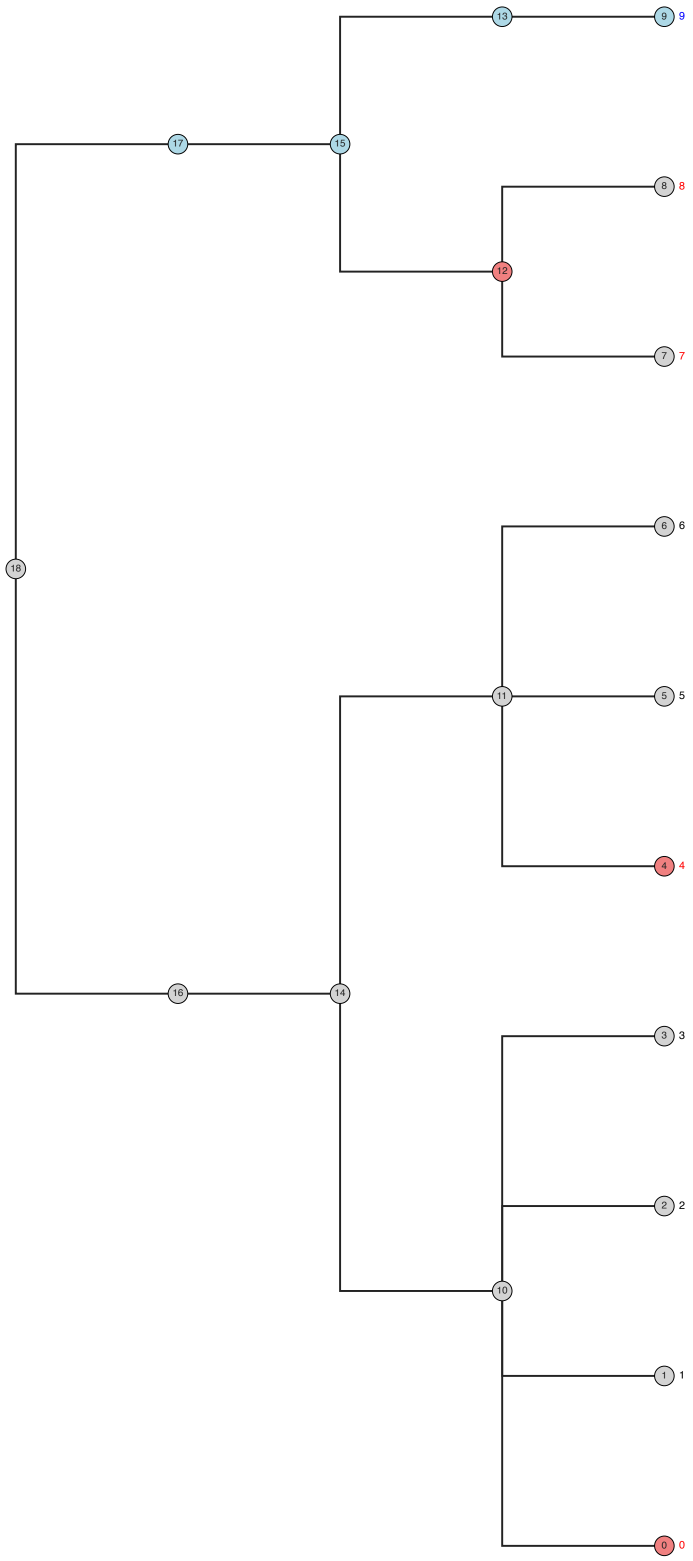


**Figure S2.** Randomly generated tree structure for synthetic data benchmark, $p = 10$ tips. The red nodes were selected to be affected by the condition, causing the red tips to be differentially abundant. The blue tip is the reference feature, which forces the effects on all blue nodes to be 0.

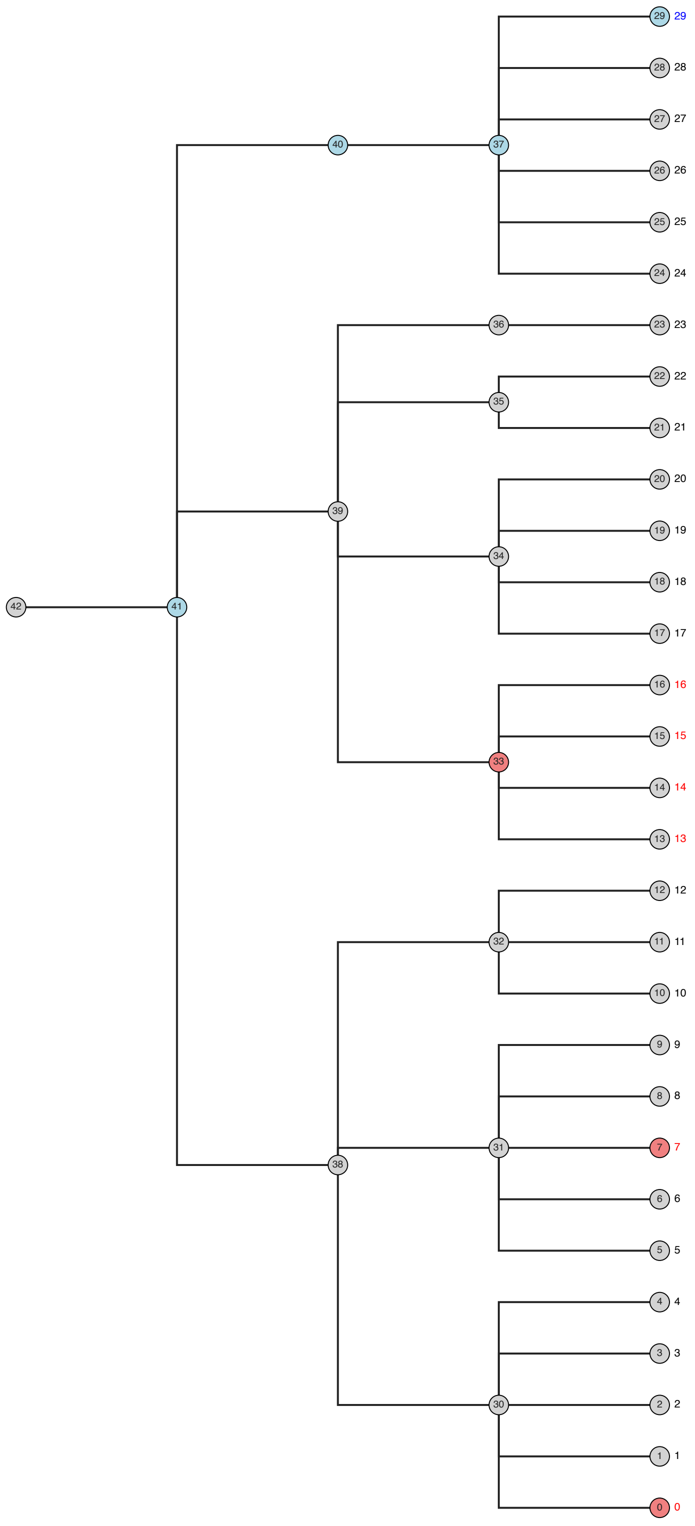


**Figure S3.** Randomly generated tree structure for synthetic data benchmark, $p = 30$ tips. The red nodes were selected to be affected by the condition, causing the red tips to be differentially abundant. The blue tip is the reference feature, which forces the effects on all blue nodes to be 0.

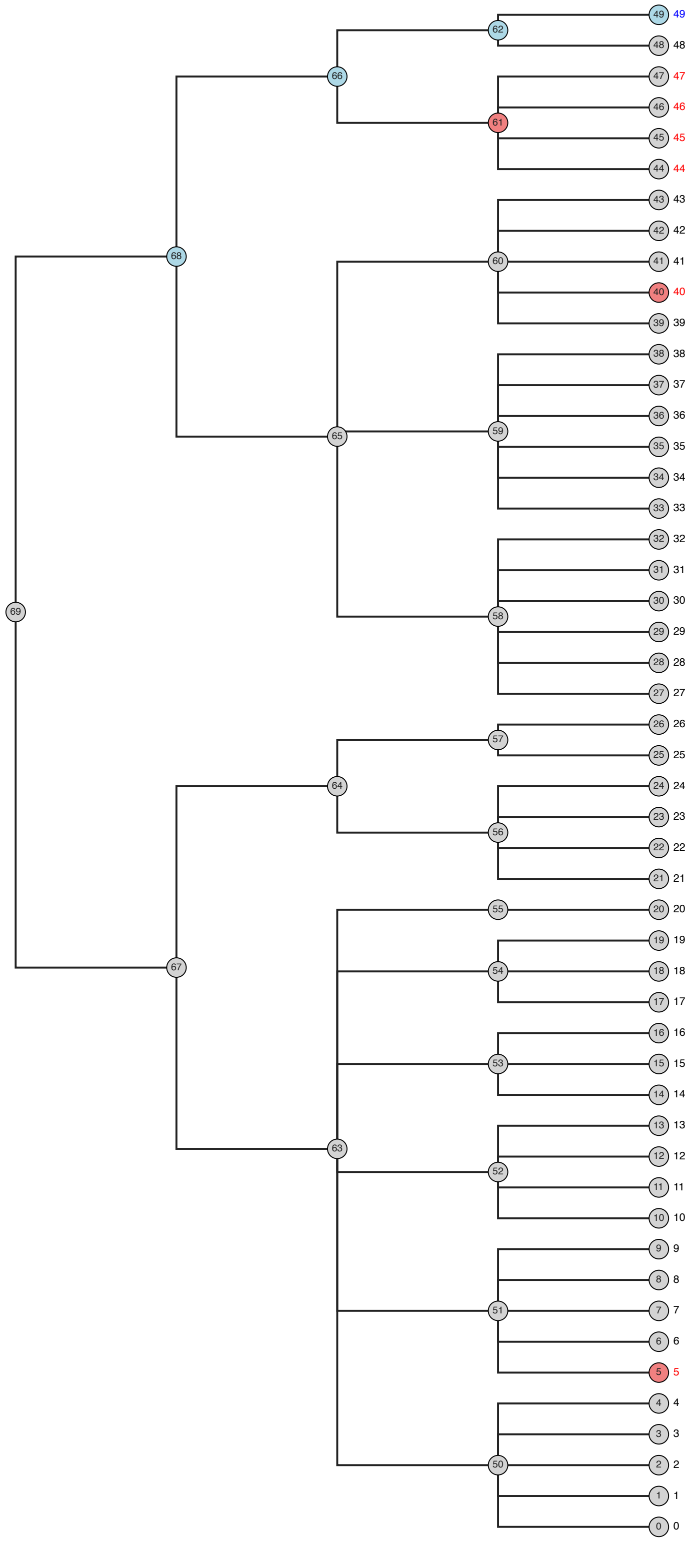


**Figure S4.** Randomly generated tree structure for synthetic data benchmark, $p = 50$ tips. The red nodes were selected to be affected by the condition, causing the red tips to be differentially abundant. The blue tip is the reference feature, which forces the effects on all blue nodes to be 0.

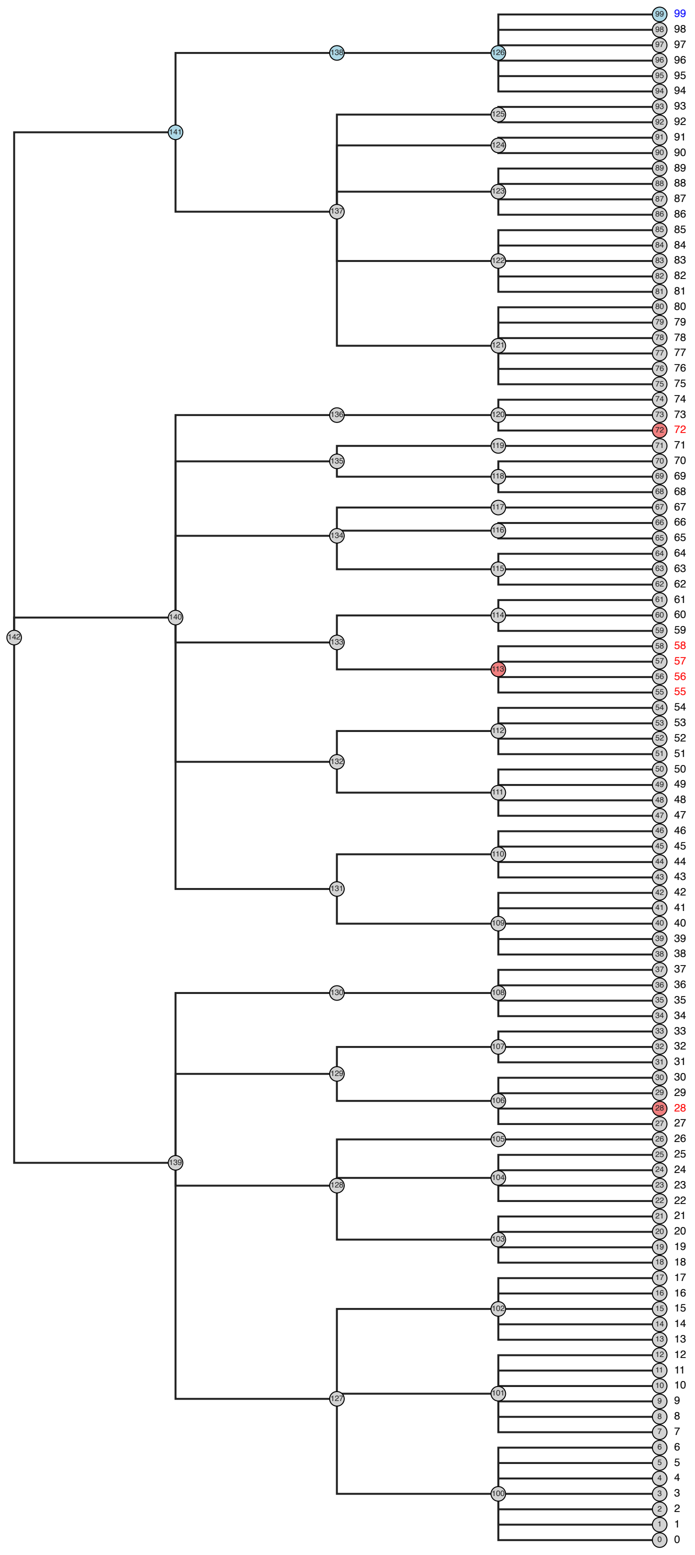


**Figure S5.** Randomly generated tree structure for synthetic data benchmark, $p = 100$ tips. The red nodes were selected to be affected by the condition, causing the red tips to be differentially abundant. The blue tip is the reference feature, which forces the effects on all blue nodes to be 0.

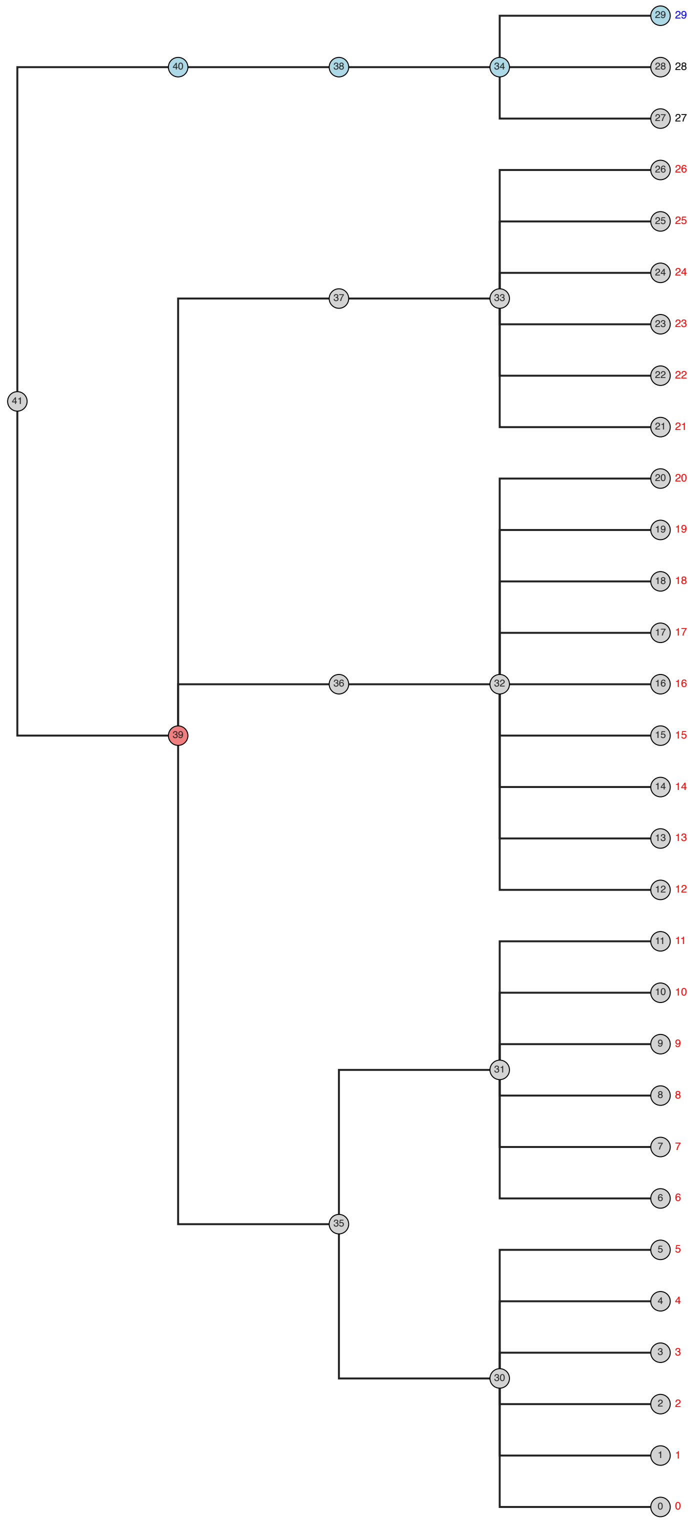


**Figure S6.** Randomly generated tree structure for synthetic data benchmark with one effect near the root of the tree, $p = 30$ tips. The red nodes were selected to be affected by the condition, causing the red tips to be differentially abundant. The blue tip is the reference feature, which forces the effects on all blue nodes to be 0.

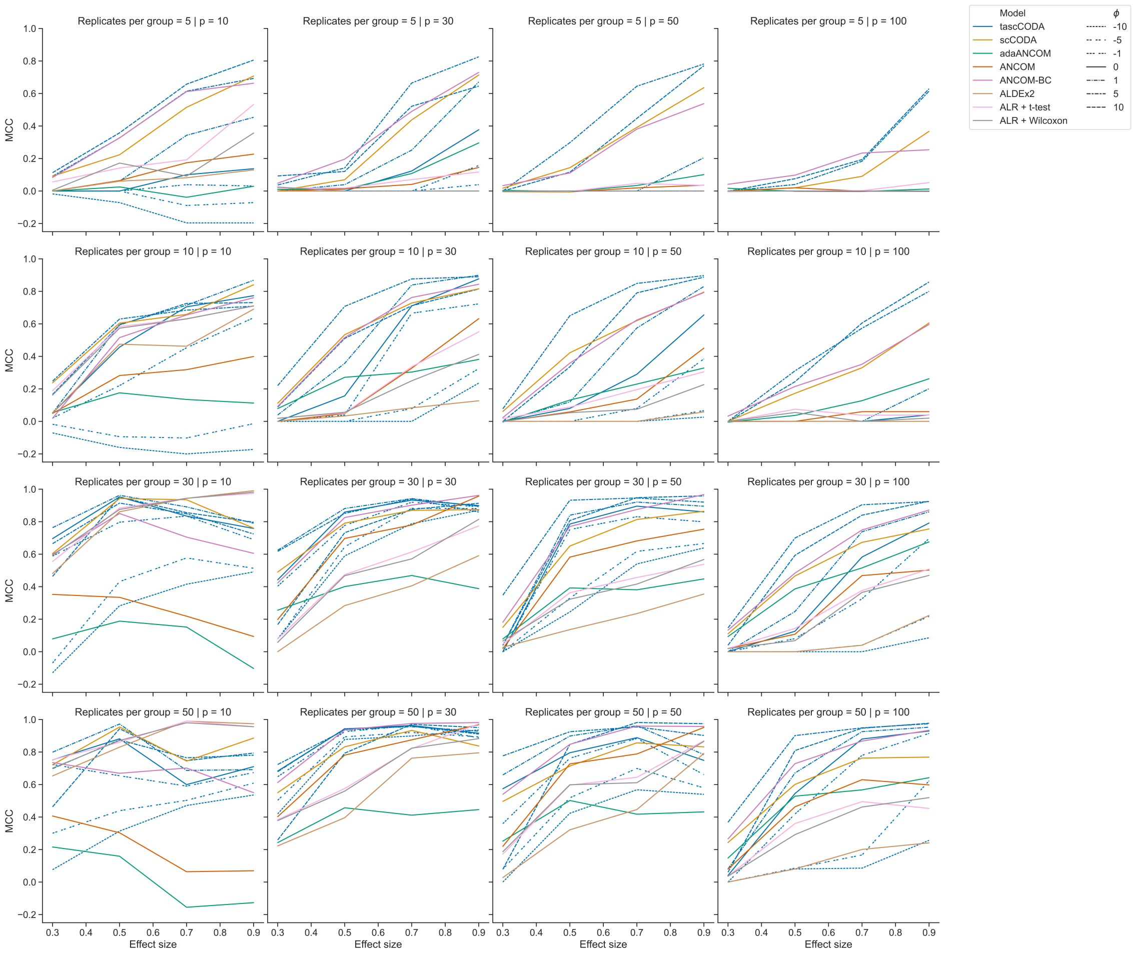


**Figure S7.** Matthews correlation coefficient (MCC) of tascCODA and other methods on simulated data with one binary covariate (differential abundance testing). Plots are grouped by the number of simulated components $p$, the number of samples per group and the effect size $\beta$. For tascCODA, different values of $\phi$ were tested (dashed blue lines).

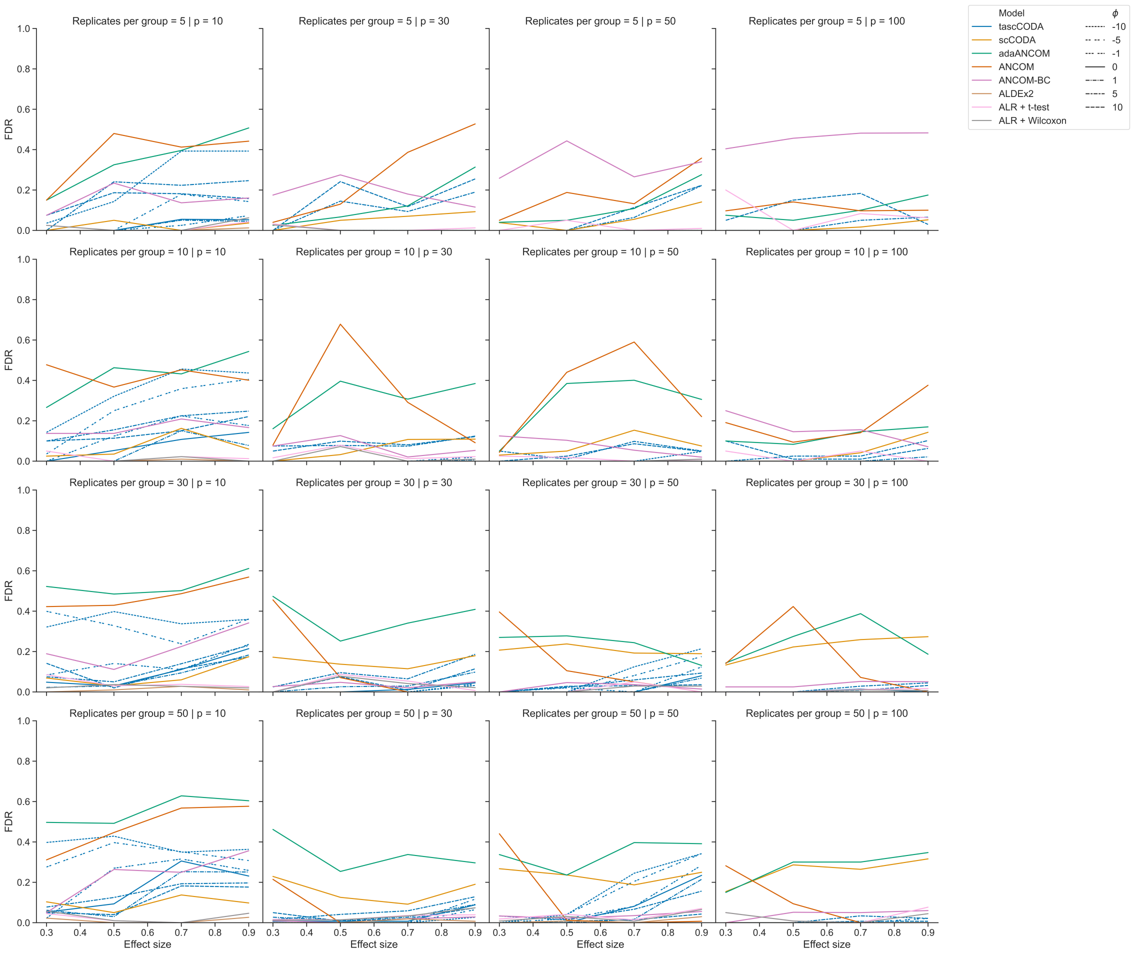


**Figure S8.** False discovery rate (FDR) of tascCODA and other methods on simulated data with one binary covariate (differential abundance testing). Plots are grouped by the number of simulated components $p$, the number of samples per group and the effect size $\beta$. For tascCODA, different values of $\phi$ were tested (dashed blue lines).

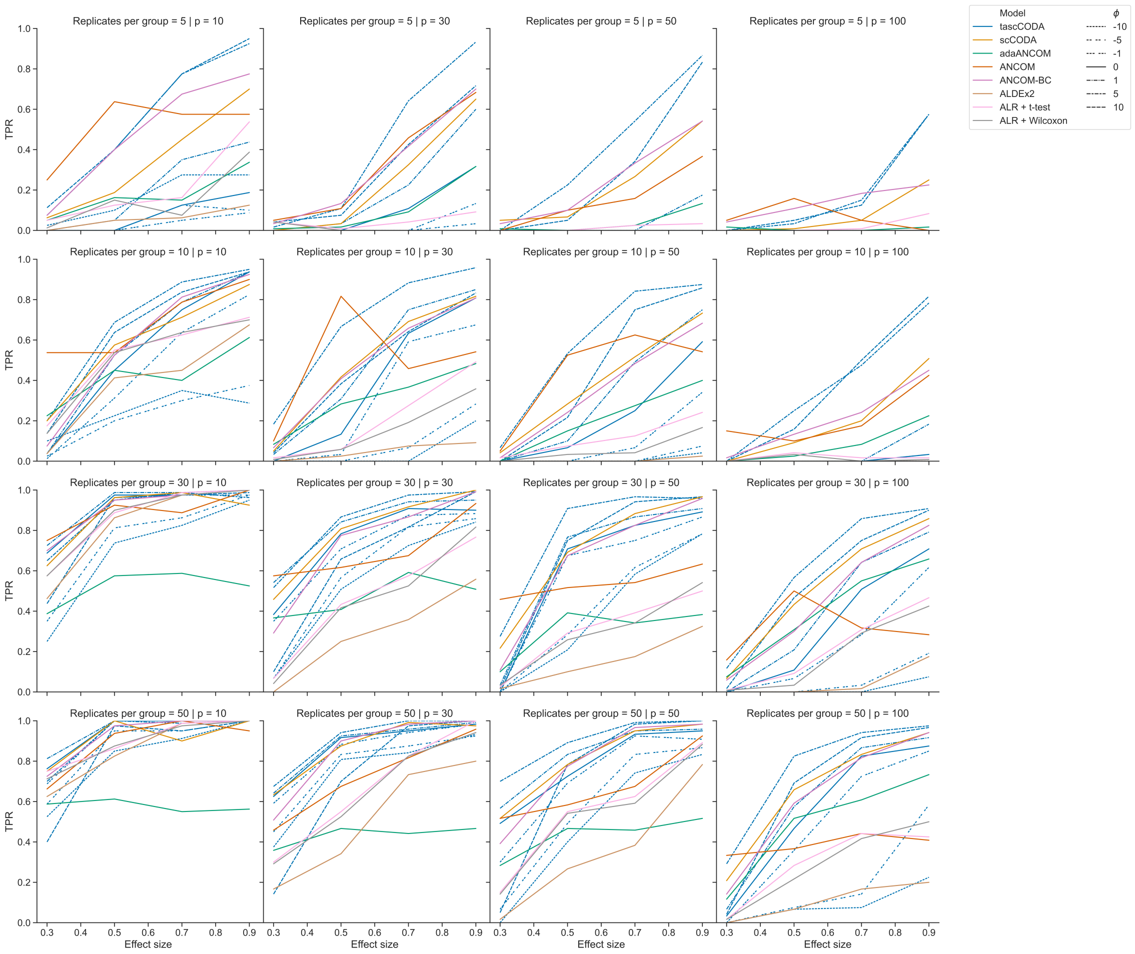


**Figure S9.** True positive rate (TPR) of tascCODA and other methods on simulated data with one binary covariate (differential abundance testing). Plots are grouped by the number of simulated components $p$, the number of samples per group and the effect size $\beta$. For tascCODA, different values of $\phi$ were tested (dashed blue lines).

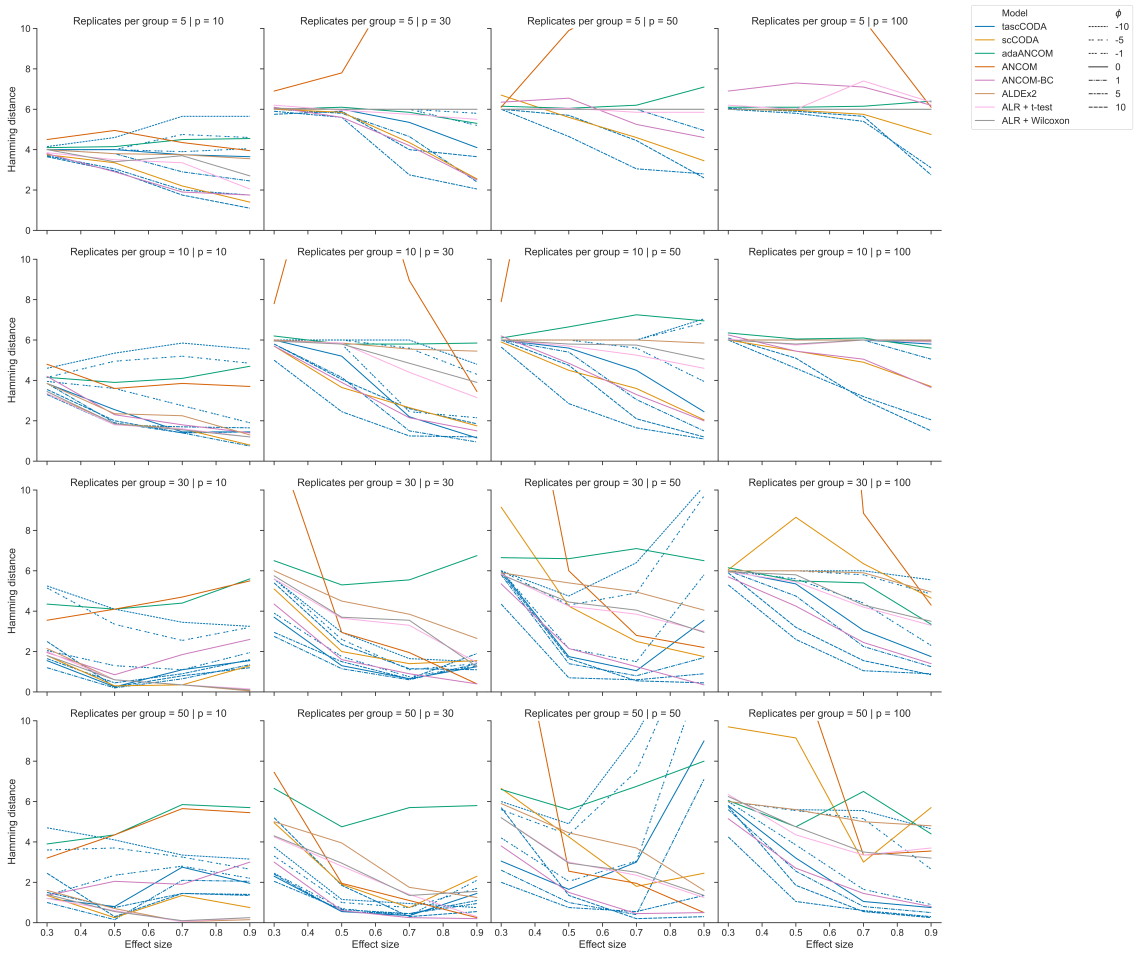


**Figure S10.** Hamming distance between ground truth and affected features determined by tascCODA and other methods on simulated data with one binary covariate (differential abundance testing). Plots are grouped by the number of simulated components $p$, the number of samples per group and the effect size $\beta$. For tascCODA, different values of $\phi$ were tested (dashed blue lines).

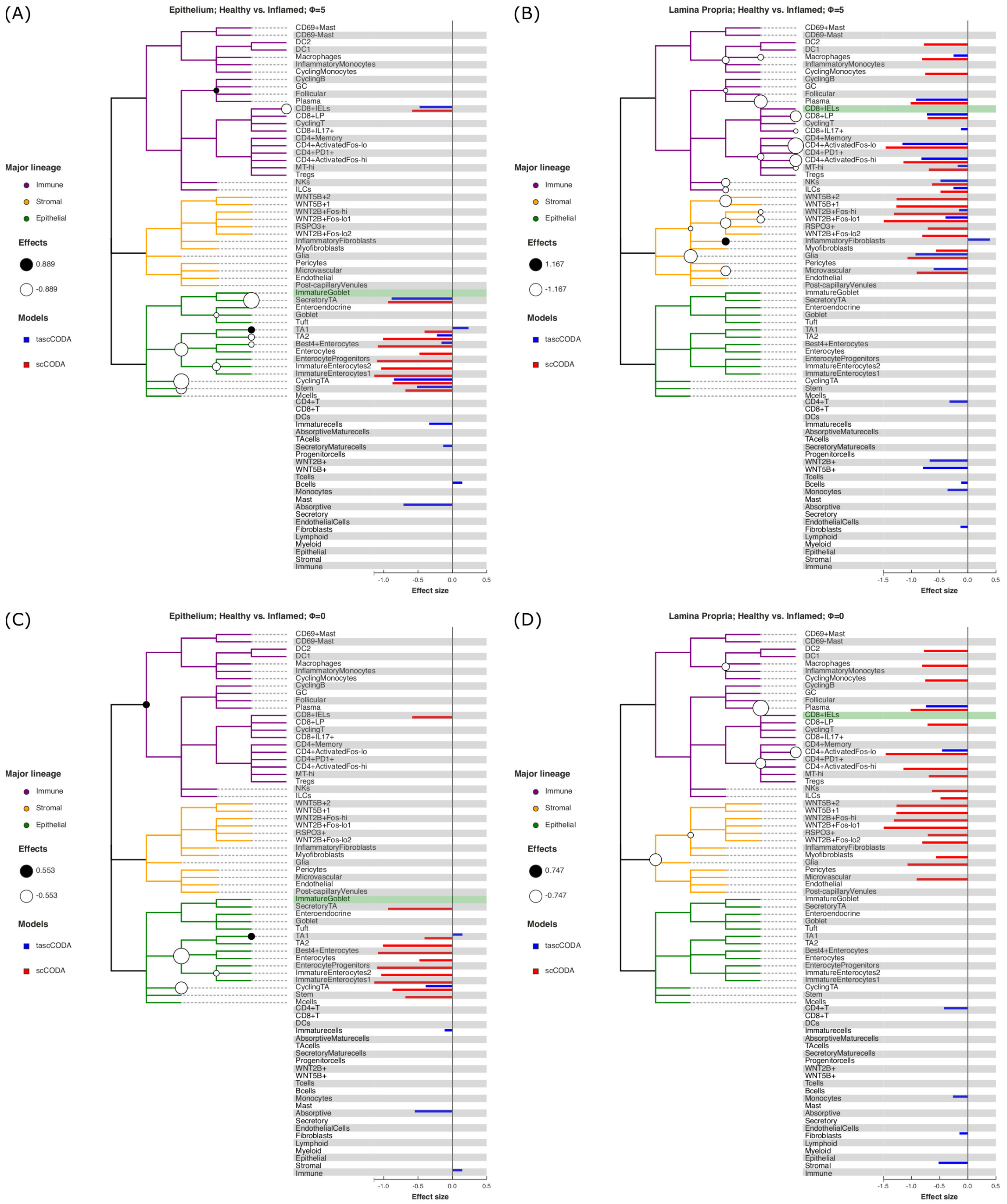


**Figure S11.** Behavior of tascCODA on scRNA-seq data. All plots show the case of comparing healthy control samples to inflamed tissue samples of UC patients in the data of Smillie et al. (2019). White and black circles on the cell lineage tree show the effects found by tascCODA, which are also shown as blue bars on the right side of each plot. The bars below the tree depict effects on internal nodes, with lower positions in the diagram corresponding to nodes closer to the root. For comparison, the red bars indicate effects found by scCODA, which only operates on the tips of the tree, on the same data. The green-shaded area shows the reference cell type that was used for both models. **(A)** $\phi = 5$, Epithelium. **(B)** $\phi = 5$, Lamina Propria. **(C)** $\phi = -3$, Epithelium. **(D)** $\phi = -3$, Lamina Propria.

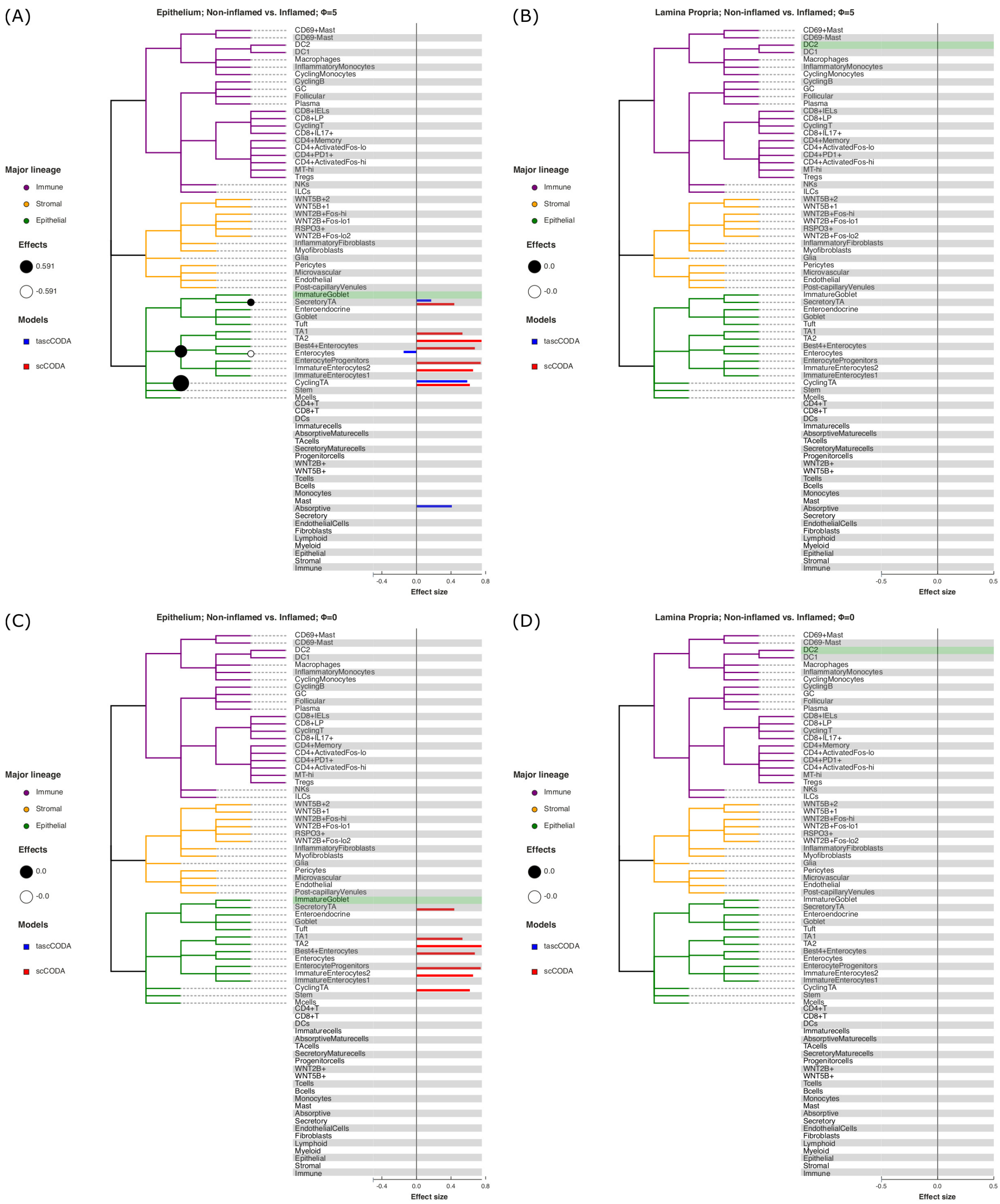


**Figure S12.** Behavior of tascCODA on scRNA-seq data. All plots show the case of non-inflamed to inflamed tissue samples of UC patients in the data of Smillie et al. (2019). White and black circles on the cell lineage tree show the effects found by tascCODA, which are also shown as blue bars on the right side of each plot. The bars below the tree depict effects on internal nodes, with lower positions in the diagram corresponding to nodes closer to the root. For comparison, the red bars indicate effects found by scCODA, which only operates on the tips of the tree, on the same data. The green-shaded area shows the reference cell type that was used for both models. **(A)** $\phi = 5$, Epithelium. **(B)** $\phi = 5$, Lamina Propria. **(C)** $\phi = -3$, Epithelium. **(D)** $\phi = -3$, Lamina Propria.

**Table S1.** Credible effects, highest density intervals, standard deviations and credibility threshold $\delta$ determined by tascCODA on epithelial biopsies from Smillie et al. (2019), $\phi = 5$. Abbreviations for scenarios: Healthy (H), Non-inflamed (N), and Inflamed (I).

| Scenario | Node | Effect | HDI 3% | HDI 97% | SD | $\delta$ |
|---|---|---|---|---|---|---|
| Epithelium - H vs. N | ImmatureEnterocytes1 | -0.647 | -0.984 | -0.309 | 0.181 | 0.148 |
| | Enterocytes | -0.211 | -0.637 | 0.054 | 0.211 | 0.148 |
| | TA1 | 0.280 | -0.022 | 0.475 | 0.134 | 0.148 |
| Epithelium - H vs. I | Stem | -0.518 | -1.000 | 0.004 | 0.286 | 0.135 |
| | CyclingTA | -0.855 | -1.144 | -0.592 | 0.146 | 0.135 |
| | Best4+Enterocytes | -0.163 | -0.893 | 0.141 | 0.303 | 0.135 |
| | TA2 | -0.229 | -0.802 | 0.129 | 0.282 | 0.135 |
| | TA1 | 0.240 | -0.159 | 0.730 | 0.265 | 0.135 |
| | SecretoryTA | -0.889 | -1.334 | -0.477 | 0.228 | 0.135 |
| | CD8+IELs | -0.481 | -1.024 | 0.018 | 0.303 | 0.135 |
| | Immaturecells | -0.343 | -0.951 | 0.091 | 0.317 | 0.132 |
| | SecretoryMaturecells | -0.138 | -0.526 | 0.082 | 0.177 | 0.132 |
| | Bcells | 0.149 | -0.068 | 0.525 | 0.178 | 0.130 |
| | Absorptive | -0.717 | -1.209 | -0.110 | 0.299 | 0.126 |
| Epithelium - N vs. I | CyclingTA | 0.591 | 0.302 | 0.907 | 0.161 | 0.144 |
| | Enterocytes | -0.152 | -0.856 | 0.121 | 0.294 | 0.144 |
| | SecretoryTA | 0.174 | -0.075 | 0.677 | 0.227 | 0.144 |
| | Absorptive | 0.413 | -0.031 | 0.742 | 0.247 | 0.135 |

**Table S2.** Credible effects, highest density intervals, standard deviations and credibility threshold $\delta$ determined by tascCODA on Lamina Propria biopsies from Smillie et al. (2019), $\phi = 5$. Abbreviations for scenarios: Healthy (H), Non-inflamed (N), and Inflamed (I). For the N vs. I scenario, no credible effects were found.

| Scenario | Node | Final Parameter | HDI 3% | HDI 97% | SD | Delta |
|---|---|---|---|---|---|---|
| LP - H vs. N | ImmatureGoblet | 0.154 | -0.161 | 0.787 | 0.270 | 0.131 |
| | Microvascular | -0.354 | -0.899 | 0.089 | 0.292 | 0.131 |
| | Glia | -0.351 | -0.867 | 0.083 | 0.278 | 0.131 |
| | ILCs | -0.189 | -0.710 | 0.125 | 0.242 | 0.131 |
| | CD4+ActivatedFos-hi | -0.144 | -0.544 | 0.100 | 0.184 | 0.131 |
| | CD4+ActivatedFos-lo | -0.608 | -1.048 | -0.162 | 0.233 | 0.131 |
| | CD8+LP | -0.169 | -0.655 | 0.105 | 0.220 | 0.131 |
| | Plasma | -0.472 | -0.895 | 0.006 | 0.238 | 0.131 |
| | TAcells | 0.469 | -0.038 | 0.952 | 0.281 | 0.129 |
| | WNT2B+ | -0.402 | -0.772 | 0.043 | 0.245 | 0.126 |
| | WNT5B+ | -0.458 | -0.935 | 0.067 | 0.296 | 0.129 |
| | Tcells | -0.438 | -0.778 | 0.043 | 0.263 | 0.117 |
| | Bcells | -0.601 | -1.047 | -0.163 | 0.229 | 0.126 |
| | Monocytes | -0.421 | -0.817 | 0.044 | 0.258 | 0.124 |
| LP - H vs. I | Microvascular | -0.612 | -1.188 | 0.040 | 0.352 | 0.126 |
| | Glia | -0.935 | -1.558 | -0.240 | 0.341 | 0.126 |
| | InflammatoryFibroblasts | 0.397 | -0.155 | 1.425 | 0.481 | 0.126 |
| | WNT2B+Fos-lo1 | -0.403 | -1.030 | 0.097 | 0.332 | 0.126 |
| | WNT2B+Fos-hi | -0.160 | -0.838 | 0.165 | 0.292 | 0.126 |
| | ILCs | -0.261 | -0.820 | 0.104 | 0.272 | 0.126 |
| | NKs | -0.491 | -0.964 | 0.025 | 0.277 | 0.126 |
| | MT-hi | -0.186 | -0.813 | 0.170 | 0.280 | 0.126 |
| | CD4+ActivatedFos-hi | -0.830 | -1.333 | -0.310 | 0.272 | 0.126 |
| | CD4+ActivatedFos-lo | -1.167 | -1.686 | -0.654 | 0.276 | 0.126 |
| | CD8+IL17+ | -0.127 | -0.685 | 0.129 | 0.237 | 0.126 |
| | CD8+LP | -0.732 | -1.044 | -0.409 | 0.168 | 0.126 |
| | Plasma | -0.925 | -1.217 | -0.580 | 0.174 | 0.126 |
| | Macrophages | -0.259 | -0.811 | 0.097 | 0.266 | 0.126 |
| | CD4+T | -0.331 | -0.693 | 0.027 | 0.213 | 0.118 |
| | WNT2B+ | -0.683 | -1.360 | 0.073 | 0.453 | 0.122 |
| | WNT5B+ | -0.803 | -1.551 | 0.051 | 0.489 | 0.125 |
| | Bcells | -0.125 | -0.474 | 0.089 | 0.162 | 0.122 |
| | Monocytes | -0.365 | -0.848 | 0.068 | 0.283 | 0.120 |
| | Fibroblasts | -0.136 | -1.052 | 0.125 | 0.362 | 0.116 |

**Table S3.** Credible effects, highest density intervals, standard deviations and credibility threshold $\delta$ determined by tascCODA on biopsies from Smillie et al. (2019), $\phi = 0$. Abbreviations for scenarios: Healthy (H), Non-inflamed (N), and Inflamed (I). Credible effects were only found for one of six scenarios.

| Scenario | Node | Effect | HDI 3% | HDI 97% | SD | $\delta$ |
|---|---|---|---|---|---|---|
| Epithelium - H vs. I | CyclingTA | -0.394 | -0.669 | 0.010 | 0.193 | 0.074 |
| | TA1 | 0.151 | -0.023 | 0.496 | 0.176 | 0.074 |
| | Immaturecells | -0.117 | -0.500 | 0.026 | 0.177 | 0.074 |
| | Absorptive | -0.553 | -0.853 | -0.205 | 0.179 | 0.074 |
| | Immune | 0.149 | -0.015 | 0.324 | 0.108 | 0.074 |
| LP - H vs. N | Plasma | -0.086 | -0.524 | 0.037 | 0.185 | 0.066 |
| | Tcells | -0.612 | -0.796 | -0.425 | 0.100 | 0.066 |
| | Bcells | -0.761 | -1.011 | -0.380 | 0.173 | 0.066 |
| | Monocytes | -0.315 | -0.618 | 0.024 | 0.216 | 0.066 |
| | Myeloid | -0.113 | -0.511 | 0.035 | 0.184 | 0.066 |
| | Epithelial | 0.145 | -0.013 | 0.322 | 0.106 | 0.066 |
| | Stromal | -0.303 | -0.483 | 0.007 | 0.143 | 0.066 |
| LP - H vs. I | CD4+ActivatedFos-lo | -0.463 | -0.967 | 0.034 | 0.316 | 0.063 |
| | Plasma | -0.747 | -0.963 | -0.528 | 0.117 | 0.063 |
| | CD4+T | -0.425 | -0.708 | -0.055 | 0.164 | 0.063 |
| | Monocytes | -0.269 | -0.568 | 0.019 | 0.197 | 0.063 |
| | Fibroblasts | -0.154 | -0.638 | 0.038 | 0.222 | 0.063 |
| | Stromal | -0.525 | -0.835 | -0.148 | 0.184 | 0.063 |

**Table S4.** Credible effects found by tascCODA comparing the gut microbiome of healthy controls and IBS patients from Labus et al. (2017) for varying aggregation levels $\phi$.

| $\phi$ | Kingdom | Phylum | Class | Order | Family | Genus | Effect |
|---|---|---|---|---|---|---|---|
| -5 | Bacteria | Firmicutes | | | | | -0.313 |
| 0 | Bacteria | Bacteroidota | Bacteroidia | Bacteroidales | Tannerellaceae | Parabacteroides | -0.156 |
| 0 | Bacteria | Bacteroidota | Bacteroidia | Bacteroidales | Bacteroidaceae | Bacteroides | -0.662 |
| 0 | Bacteria | Firmicutes | Clostridia | Oscillospirales | | | -0.232 |
| 5 | Bacteria | Bacteroidota | Bacteroidia | Bacteroidales | Tannerellaceae | Parabacteroides | -0.845 |
| 5 | Bacteria | Bacteroidota | Bacteroidia | Bacteroidales | Bacteroidaceae | Bacteroides | -1.001 |
| 5 | Bacteria | Bacteroidota | Bacteroidia | Bacteroidales | Prevotellaceae | Prevotella | -0.413 |
| 5 | Bacteria | Firmicutes | Clostridia | Lachnospirales | Lachnospiraceae | Agathobacter | -0.610 |
| 5 | Bacteria | Firmicutes | Clostridia | Oscillospirales | Ruminococcaceae | Subdoligranulum | -0.224 |
| 5 | Bacteria | Firmicutes | Clostridia | Oscillospirales | Ruminococcaceae | Faecalibacterium | -0.252 |
| 5 | Bacteria | Firmicutes | Negativicutes | Acidaminococcales | Acidaminococcaceae | Phascolarctobacterium | -0.250 |
| 5 | Bacteria | Firmicutes | Clostridia | Oscillospirales | Ruminococcaceae | | -0.340 |

**Table S5.** Credible effects found by tascCODA ($\phi = 5$) comparing the gut microbiome of four different subtypes of IBS to all other samples. Original data by Labus et al. (2017).

| Subtype | Kingdom | Phylum | Class | Order | Family | Genus | Effect |
|---|---|---|---|---|---|---|---|
| IBS-C | Bacteria | Bacteroidota | Bacteroidia | Bacteroidales | Bacteroidaceae | Bacteroides | -0.426 |
| IBS-C | Bacteria | Firmicutes | Clostridia | Lachnospirales | Lachnospiraceae | Anaerostipes | 0.438 |
| IBS-C | Bacteria | Firmicutes | Clostridia | Lachnospirales | Lachnospiraceae | Agathobacter | -0.819 |
| IBS-C | Bacteria | Firmicutes | Clostridia | Oscillospirales | Ruminococcaceae | Ruminococcus | -0.262 |
| IBS-C | Bacteria | Firmicutes | Clostridia | Oscillospirales | Ruminococcaceae | Faecalibacterium | -0.320 |
| IBS-D | Bacteria | Bacteroidota | Bacteroidia | Bacteroidales | Tannerellaceae | Parabacteroides | -0.392 |
| IBS-D | Bacteria | Bacteroidota | Bacteroidia | Bacteroidales | Bacteroidaceae | Bacteroides | -1.405 |
| IBS-M | Bacteria | Bacteroidota | Bacteroidia | Bacteroidales | Tannerellaceae | Parabacteroides | -0.424 |
| IBS-M | Bacteria | Firmicutes | Clostridia | Lachnospirales | Lachnospiraceae | Blautia | 0.799 |
| IBS-M | Bacteria | Firmicutes | Clostridia | Oscillospirales | Ruminococcaceae | Faecalibacterium | -0.285 |
| IBS-unspecified | Bacteria | Firmicutes | Clostridia | Peptostreptococcales-Tissierellales | Peptostreptococcaceae | Romboutsia | 0.259 |
